# *Musashi* expression in intestinal stem cells attenuates radiation-induced decline in intestinal homeostasis and survival in *Drosophila*

**DOI:** 10.1101/540377

**Authors:** Amit Sharma, Kazutaka Akagi, Blaine Pattavina, Kenneth A Wilson, Christopher Nelson, Mark Watson, Elie Maksoud, Mauricio Ortega, Rachel Brem, Pankaj Kapahi

**Affiliations:** Buck Institute for Research on Aging, 8001 Redwood Boulevard, Novato, CA 94945, USA; National Center for Geriatrics and Gerontology, 7-430 Morioka-cho, Obu, Aichi 474-8511, Japan

## Abstract

Exposure to genotoxic stress by environmental agents or treatments, such as radiation therapy, can diminish health span and accelerate aging. We have developed a *Drosophila melanogaster* model to study the molecular effects of radiation-induced damage and repair. Utilizing a quantitative intestinal permeability assay, we performed an unbiased GWAS screen (using ∼150 strains from the DGRP reference panel) to search for natural genetic variants that regulate radiation-induced gut permeability. From this screen, we identified the RNA binding protein, *Musashi (msi).Msi* overexpression promoted intestinal stem cell proliferation, which increased survival after irradiation and rescued radiation-induced gut permeability. We identified that a novel role of *Msi in* ISC proliferation a novel role for *Msi* in enhancing ISC function following radiation-induced gut damage, which identifies *Msi* as a potential therapeutic target.

## Introduction

A typical mammalian cell encounters approximately 2 × 10^5^ DNA lesions per day [1]. External stressors, both chemical and radioactive, and internal factors, such as oxidative stress, are the primary sources of DNA damage [2]. The inability to correct DNA damage results in the accumulation of harmful mutations, which contribute to cellular damage, cancer, and aging [3–8]. However, DNA damaging agents, such as radiation, are the only available treatments for certain pathologies. These therapies can lead to complications due to cellular and tissue damage caused by genotoxic stress. For example, genetic and epigenetic alterations in the tumor [9], or tumor microenvironment, may render it resistant to radiation [10]. Additionally, bystander tissues can be damaged from radiotherapy [11, 12]. Patients undergoing radiotherapy encounter both short-term side effects (nausea, vomiting, and damage to epithelial surfaces) and long-term side effects (enteritis [13], radiation proctitis, heart disease [14], and cognitive decline [15]).

Organisms have developed several DNA error correction mechanisms, as the inability to correct DNA errors leads to permanent cellular damage [16–19]. To prevent the propagation of DNA damage and improve survival following iradiation, cells choose from among several fates: cells may apoptose [26], enter a state of replicative arrest, such as senescence, or be cleared by phagocytosis or autophagy. [20]. However, *in vitro* models of genotoxic stress fail to recapitulate these fates; they fail to represent the complexities of tissue microenvironments and the cell-nonautonomous consequences of radiation damage. For instance, apoptotic or senescent cells may produce secreted factors that exacerbate damage to cells that did not receive the primary insult [21].

As the gastrointestinal tract encompasses a large area in the body, it is commonly a bystander tissue in radiotherapy and accounts for a significant percentage of side effects from radiation treatment [22, 23]. The fly and human intestines share similar tissue, anatomy, and physiological function [24, 25]; both fly and mammalian guts are composed of ISCs, enterocytes, and enteroendocrine (EE) cells [26]. Intestinal stem cells (ISCs) are involved in regenerative and tissue-repair processes [27, 28] in flies and mammals [29]. DNA damage to the ISCs leads to a reduced proliferative potential, which contributes to the pathogenesis of radiation enteritis in patients undergoing radiation therapy. [30–33].Previous studies have used *D. melanogaster* to identify conserved molecular pathways that maintain stem cells’ physiological function, tissue repair, and homeostasis in the gut [34–36]. Here, we have taken advantage of the flies’ genetic malleability, short lifespan, and complex tissue microenvironments to develop a whole-animal model to study therapeutic targets for radiation damage to the intestine.

We show that radiation significantly damages flies’ intestine, which decreases ISC proliferation and survival, and increases intestinal permeability and inflammation. We use the ***Drosophila* Genetic Reference Panel (DGRP),** a collection of approximately 150 fully sequenced fly strains, as a screening tool to identify natural variants that regulate radiation-induced intestinal permeability [37]. One candidate gene we validate is the RNA binding protein, **Musashi (msi)**, which regulates the expression of target genes by binding to their 3’UTR using a consensus sequence [38]. Msi is also involved in neuronal differentiation and cell fate determination [39, 40]. Here we demonstrate that ISC-specific knockdown of *msi* significantly increases gut permeability after irradiation. Conversely, we find that overexpressing *msi* reduces gut permeability and enhances survival after irradiation. Interestingly, we observe that *msi* overexpression increases ISC proliferation and tissue repair. We find that after irradiation, *Msi*-mediates ISC proliferation by indirectly regulating **cyclic AMP (cAMP)** levels through post-transcriptional modulation of Ac13e. Thus, we identify *msi* as a potential therapeutic target for maintaining gut homeostasis after exposure to radiation.

## Methodology

### Fly culture, stocks and lifespan analysis

Flies were reared on standard laboratory diet (Caltech food recipe: 8.6% Cornmeal, 1.6% Yeast, 5% Sucrose, 0.46% Agar, 1% Acid mix) [41, 42]. Emerged adults were transferred within 3-5 days to the **yeast extract (YE)** diet (1.5% YE, 8.6% Cornmeal, 5% Sucrose, 0.46% Agar, 1% Acid mix). For *Gene-Switch Gal4* drivers, RU486 was dissolved in 95% ethanol with a final concentration of 100μM (the media is then referred to as ‘+RU486’) and was administered from the adult stage (5 day old). The control diet contained the same volume of 95% ethanol and is referred to as ‘RU486’. Life spans were analyzed as described previously [41, 43].

### Radiation exposure

Adult, female, 5-day old flies were exposed to different doses of X-rays at 320 kV and 10 mA to achieve the required doses as indicated and maintained on a standard fly diet.

### Quantitative Real-time-PCR

Total RNA was extracted from 12 female guts, 8 female fat bodies (fly abdomen) or 5 female whole flies using Quick-RNA MiniPrep Kit (Zymo Research). cDNA was synthesized using QuantiTect Reverse Transcription Kit (QIAGEN). 1 µg of total RNA was used per sample. qPCR reaction was performed in duplicate on each of 3 independent biological replicates using SensiFAST SYBR No-ROX Kit (BIOLINE). Error bars indicate standard deviation. Samples were normalized with *ribosomal protein 49* (*rp49*), or *glyceraldehyde-3-phosphate dehydrogenase* (*GAPDH*).

### Immunohistochemistry

Dissected guts were fixed with 4% paraformaldehyde in PEM, 100 mM Pipes, 2 mM EGTA, 1 mM MgSO_4_, for 45 minutes. Samples were washed for 5 minutes three times with PEM, then incubated with 1% NP40/PEM for 30 minutes. Samples were washed for 10 minutes three times with TBS-TB, 0.02 % Triton X-100/PBS, 0.2 % BSA, and blocking was performed with 5% goat serum in TBS-TB for 2 hours at 25°C. Samples were incubated with the primary antibody overnight at 4°C, then washed for 10 minutes three times with TBS-TB, and incubated with the secondary antibody for 2 hours at room temperature. Nuclei were stained using DAPI. Samples were mounted with Mowiol mounting buffer and analyzed by confocal microscope (Zeiss: LSM780).

### Acridine orange staining

Dissected guts were incubated with Ethidium Bromide and acridine orange (Sigma: 5 μg/ml) (10 μg/ml) in PBS for 5 minutes at room temperature. Samples were rinsed with PBS twice, then mounted with PBS and immediately analyzed by microscope (Olympus: BX51).

### Smurf gut permeability assay

25 flies were placed in an empty vial containing a piece of 2.0 cm x 4.0 cm filter paper. 300 μl of blue dye solution, 2.5 % blue dye (FD&C #1) in 5% sucrose, was used to wet the paper as a feeding medium. Smurf and non-smurf flies were counted following incubation with feeding paper for 24 hours at 25 °C. Smurf flies were quantified as flies with any visible blue dye outside of the intestines.

### Spontaneous activity

24 hours after irradiation, flies were placed in population monitors and their physical activity was recorded every 10 minutes for 24 hours (*Drosophila* population monitor by Trikinetics Inc., Waltham, MA, USA). Reading chambers have circular rings of infrared beams at three different levels, which allow recording whenever a fly crosses the rings. Activity monitors were kept in temperature-controlled incubators set at 25°C on a 12-hour light-dark cycle. The daylight period began at 8:00 AM.

### Screening for variants associated with regulating irradiation-induced phenotypes

Two weeks following irradiation, we observed significant variation in the gut permeability assay between DGRP lines in the proportion of Smurf flies. Candidates with a false detection rate (FDR) of 27% or less were considered for further validation [37]. FDRs were calculated empirically from permuted data [44]. The association was determined by aligning phenotypic values at an allelic marker. Genetic markers with >25% minor allele frequency were used [45] by employing custom scripts written in Python, using ordinary least squares regression from the stats models module. The analysis was done using linear model: phenotype = **β**_1_xGenotype + **β**_2_xIrradiationDose+ **β**_3_xGenotype X-Irradiation Dose+ intercept.

The p-values shown reflect whether the **β** term is 0. The Genotype X-Irradiation Dose term reflects the Irradiation-dependent portion of genetic influence on the phenotype [45].

## Results

### Ionizing radiation reduces survival and locomotion in *D. melanogaster*

In the present study, we investigated the effects of irradiation on adult flies. Even though ionizing radiation (IR) has been extensively studied in the context of mutagenesis experiments [46, 47] and embryonic development signals [48] in *D. melanogaster,* not much is known regarding its effects in adult flies. We exposed 5-day old *w*^*1118*^ adult flies exposure to different doses of IR. Interestingly, these flies were fairly resistant to lower doses of X-rays (from 100R to 1000R), likely because most tissues in the fly are post-mitotic [49]. However, when we exposed female *w*^*1118*^ flies to 10kR, it significantly reduced their mean lifespan, compared to un-irradiated controls (**Fig. 1A**) (**Supplemental Fig. 1A**). We observed a similar reduction in lifespan in irradiated male flies, indicating that adult sensitivity to IR is sex independent (**Supplemental Fig. 1C**).

**Figure 1:**
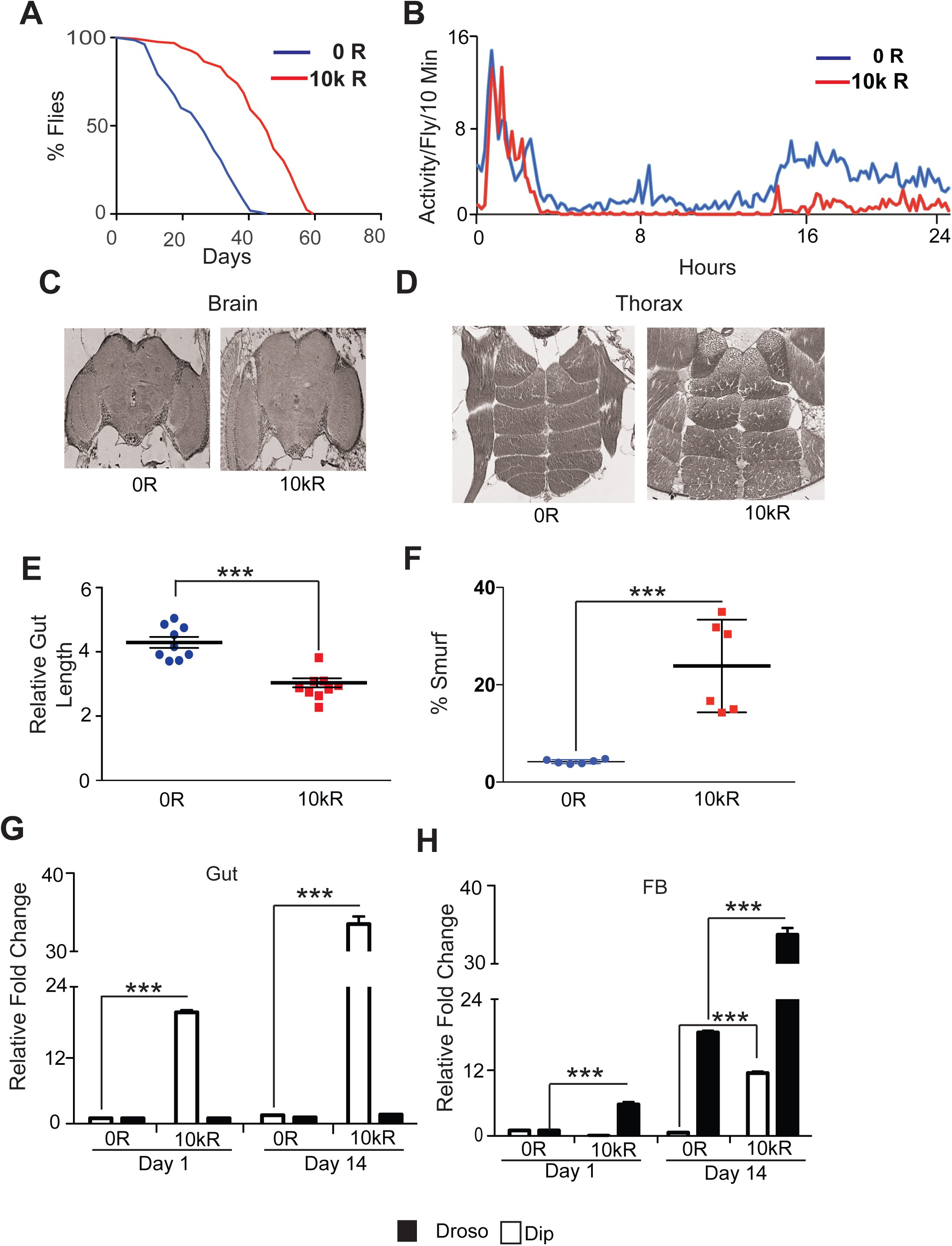
Radiation induced damage results in significant reduction in survival and spontaneous activity due to damage to the gut. **(A)** Kaplan Meier survival analysis for survival irradiation of 5 day old flies. At least 150 flies were used for both irradiated group (10k R) and non-irradiated control (0 R). **(B)** The effect of radiation on spontaneous activity. The graph shows averaged activity (four vials per group with 25 flies in each vial) per 10 minutes for control and irradiated flies. The X-axis represents time (in hrs) after the flies were moved to the activity monitors. The activity measurement was started at 4:00pm. **(C)** Representative H and E (Hematoxylin and eosin) staining of the paraffin embedded brain and **(D)** thorax of *w*^*1118*^, 7 days after irradiation showing no structural abnormalities. **(E)** The graph represents intestine length measured using ImageJ in dissected 14 days after irradiation. The relative length of the *w*^*1118*^ 5 days old adult female flies 14 days after irradiating with or without 10k R is plotted as arbitrary units. **(F)** The effect of radiation on gut permeability. Smurf assay to access gut permeability was performed in *w*^*1118*^ 5 days old adult female flies 14 days after irradiating with or without 10k R. Results were plotted as mean percentage of ’Smurf’ to non-smurf flies. Error bars indicate S.D. of 4 replicates. (** p < 0.01, * p < 0.05 by *t*-test). **(G)** Real time PCR performed with RNA prepared using dissected tissue. Results plotted as relative fold change in the expression of immune response genes *Diptericin* (*Dip*) and Drosomycin (Droso) in fat body FB and **(H)** gut. The results are present as mean relative fold change in the gene expression and normalized to housekeeping gene RP49 on days 1 and 14 after irradiation (10k R), demonstrating local and systemic inflammation.

A frequent adverse effect of radiation exposure is fatigue [50–55]. Several groups have observed that irradiated mice have diminished spontaneous and voluntary activity [56, 57]. We used the *Drosophila* Activity Monitor System to investigated whether radiation also reduces flies’ spontaneous physical activity [58]. Our results show that irradiated flies displayed both an acute (24h post-exposure to IR) and chronic (14d post-exposure to IR) reduction in spontaneous physical activity (**Fig. 1B** and **Supplementary Fig 1B**). This might be due to muscle damage however, when we evaluated the morphological changes in flight and thoracic muscles in irradiated flies, by H and E staining, we did not observe any overt structural damage to the muscles in the thorax (day one and seven after irradiation) (**Fig. 1C**). In addition, we did not observe significant structural damage to the brain, one and seven days after irradiation (not shown and **Fig. 1D**). These results indicate that neither muscle nor brain damage accounts for the reduction in survival and activity in irradiated flies.

### Ionizing radiation disrupts intestinal integrity and increases inflammation

Because disruption to flies’ intestine is known to impact survival [59], we examined whether the reduced survival of irradiated flies was from damage to intestinal tissue. Interestingly, our results showed that irradiated flies have significantly shorter intestines than non-irradiated controls 14 days after irradiation, which suggests that irradiation may structurally damage the fly’s gut (**Fig. 1D & Supplementary Fig. 1B-ii**). We hypothesized that this structural damage to the gut influences the gut’s barrier function. To test this, we measured the effect of irradiation on intestinal permeability, by performing the well-established quantitative Smurf assay [60]. Our results in **Fig. 1E** demonstrate a significantly higher percentage of flies with permeable intestine (Smurf flies) upon irradiation when compared to un-irradiated controls (14 days after irradiation). This effect of ionizing radiation on intestinal permeability was dose-dependent; increasing dosage of radaition progressivly increased the percentage of flies with permeable intestine (**Supplementary Fig. 1D**). Furthermore, radiation also induced intestinal permeability when the dosage was staggered (4 doses of 2500 R every other day) (**Supplementary Fig. 1E**). The detrimental effect of IR was sex independent, as irradiated male flies also lived significantly shorter (**Supplementary Fig. 1C**) and had increased intestinal permeability (14 days after irradiation) (**Supplementary Fig. 1F**).

The disruption of gut barrier integrity after irradiation, as indicated by the Smurf assay, may result in increased local and systemic inflammation, indicated by the secretion of anti-microbial peptides (AMPs) [60]. To test this, we investigated the effect of radiation on the expression of the sentinal AMPs, *Drosomycin (Droso)* and *Diptericin (Dip),* [61] in dissected guts and fat bodies. This served as a proxy for local (intestine) and systemic inflammation. Quantitative realtime PCR (qRT-PCR) on dissected intestinal tissue samples indicated a 20-fold increase in *Dip* expression as early as 24 hours after irradiation, which increased to 30-fold 14 days after irradiation. This coincided with the intestinal permeability observed in our Smurf data, which indicates elevated IMD signaling (Immune Deficiency) [62], a critical response to bacterial infection (**Fig.1G**). Interestingly we did not observe an appriciable increase in Drosomycin expression in the irradiated intestines.

Fat body in *Drosophila* contributes to the humoral immune response, hence their AMP production may serve as a proxy for systemic inflammation [63, 64]. Quantitative RT-PCR on samples from the fat body revealed a 4-fold increase in the expression of *Drosomycin* 1, followed by a 35-fold increase 14 days after irradiation (**Fig. 1 H**). This indicates a sustained increase in systemic inflammation from elivated Toll signaling [65] in the fat bodies. Together, these results demonstrate that irradiation induces a sustained local and systemic inflammatory response in the adult fly.

### Exposure to radiation causes DNA damage, cell death in enterocytes and inhibition of ISC proliferation

Exposure to ionizing radiation induces DNA double strand breaks (DSB) [66, 67]. One of the earliest events following DSBs is activation of kinases like ATM, ATR and DNK-PK, which phosphorylate the C-terminal tail of the histone 2A [68]. This DSB-induced phosphorylation is conserved in *Drosophila* [69, 70]. We tested the effect of ionizing radiation on histone H2Av phosphorylation in flies’ intestinal tissue by Immunofluorescence staining for γ-H2Av. Following irradiation, flies’ intestine showed a substantial increase in γ-H2Xv foci compared to non-irradiated flies (**Fig. 2A**) in a dose-dependent manner (**Supplementary Fig. 2A**). Environmental stress on gut enterocytes is known to activate reparative responses, often initiated by the IL-6-like cytokine, Upd3 [71]. We tested if persistent DNA damage, caused by radiation, affects Upd3 expression. Our results indicate that Upd3 expression is significantly upregulated 24 hours after irradiation, based on GFP reporter expression (**Fig. 2B**), the analysis of GFP fluorescent intensity indicated a 3-fold-induction of the reporter gene (**Supplementary Fig. 2B**). These results were supported by RT-qPCR of Upd3 in dissected guts from *w*^*1118*^ female flies, which showed an approximately three-fold increase in Upd3 expression, compared to un-irradiated controls (**Supplementary Fig. 2C**).

**Figure 2:**
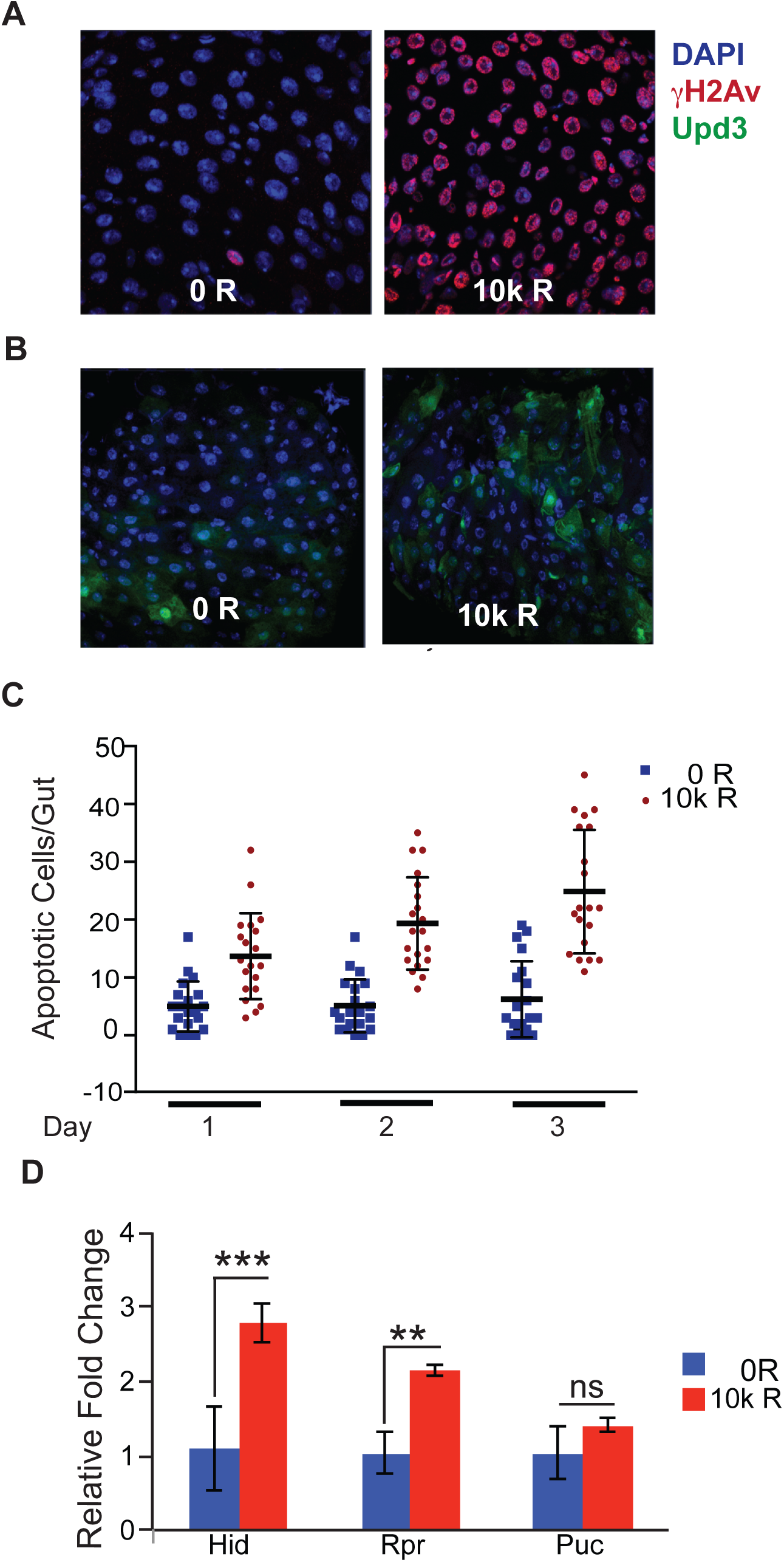
Exposure to radiation causes DNA damage and cell death in enterocytes and inhibition of ISC proliferation. **(A)** Representative image of adult *Drosophila* gut dissected 30 minutes after irradiation with or without 10k R and stained with γ-H2Av antibody (Red) and co-stained with DAPI (blue) **(B)** Representative images of the dissected gut from recombinant line, *Upd-3-Gal4 UAS-GFP*, 3 days after irradiation with or without 10k R indicates substantial up-regulation of Upd3 in ECs (green). Nuclei are counter-stained with DAPI (blue). (C) Ethidium bromide-Acridine Orange staining was performed in guts of *w*^*1118*^ female flies after irradiation (10k R) at indicated time points. The results are plotted as the mean number of apoptotic cells per gut and presented and mean apoptotic cells and error bars indicate S.D. of 2 independent experiments with at least 10 guts each. **(D)** Relative fold change in the expression of immune response genes apoptotic genes *Hid, Rpr and Puc* in the dissected gut 24 hours after irradiation. Results were plotted as mean fold change and the error bars indicate S.D. of 4 replicates. (** p < 0.01, * p < 0.05 by *t*-test).

Metazoan cells also undergo apoptosis following DNA DSB; thus we investigated the effect of radiation on apoptosis in the fly’s intestine. To this end, we performed Acridine Orange/Ethidium Bromide staining assay in dissected intestinal tissue, which showed an approximate two-fold increase in the number of apoptotic cells on days 1, 2 and 3 following irradiation [72] (**Fig. 2C**). These results were also supported by qRT-PCR (using RNA prepared from dissected intestine), which showed elevated expression of the pro-apoptotic genes, *hid, reaper* [73] and *puckered* [74] 24 hours after irradiation (**Fig. 2D**). We observed a substantially lower number of apoptotic cells in the gut of flies irradiated with lower doses of radiation (1k-5k R) (**Supplementary Fig. 2D**).

### Exposure to X-rays disrupts intestinal homeostasis and ISC proliferation

Previous studies have shown that fly guts respond to damage from toxins like DSS or Bleomycin [75], and stress from bacterial infection [76], by inducing the proliferation of ISCs (Intestinal Stem cells), which enhances intestinal repair by replacing damaged cells [77]. We investigated if ISCs in irradiated flies restore tissue homeostasis by replacing apoptotic enterocytes. To test this, we stained guts with an anti-phosphohistone H3 (anti-PH3) antibody that marks dividing cells. Immunofluorescence staining in dissected guts demonstrated that irradiation inhibited ISC proliferation as early as 1 day after irradiation **(Fig. 3A**). This inability of ISCs to repair damage was even observed 14 days after irradiation (**Fig. 3B**). We reasoned that inhibiting ISC proliferation affects ISC numbers. To test this, we irradiated a recombinant fly line carrying a Dl-LacZ enhancer trap that has been extensively used to identify ISCs [78, 79]. We observed that exposure to radiation significantly reduced the numbers of *delta* positive ISCs and ISC marker (1 and 14 days after irradiation) (**Fig. 3C & 3D**).

**Figure 3:**
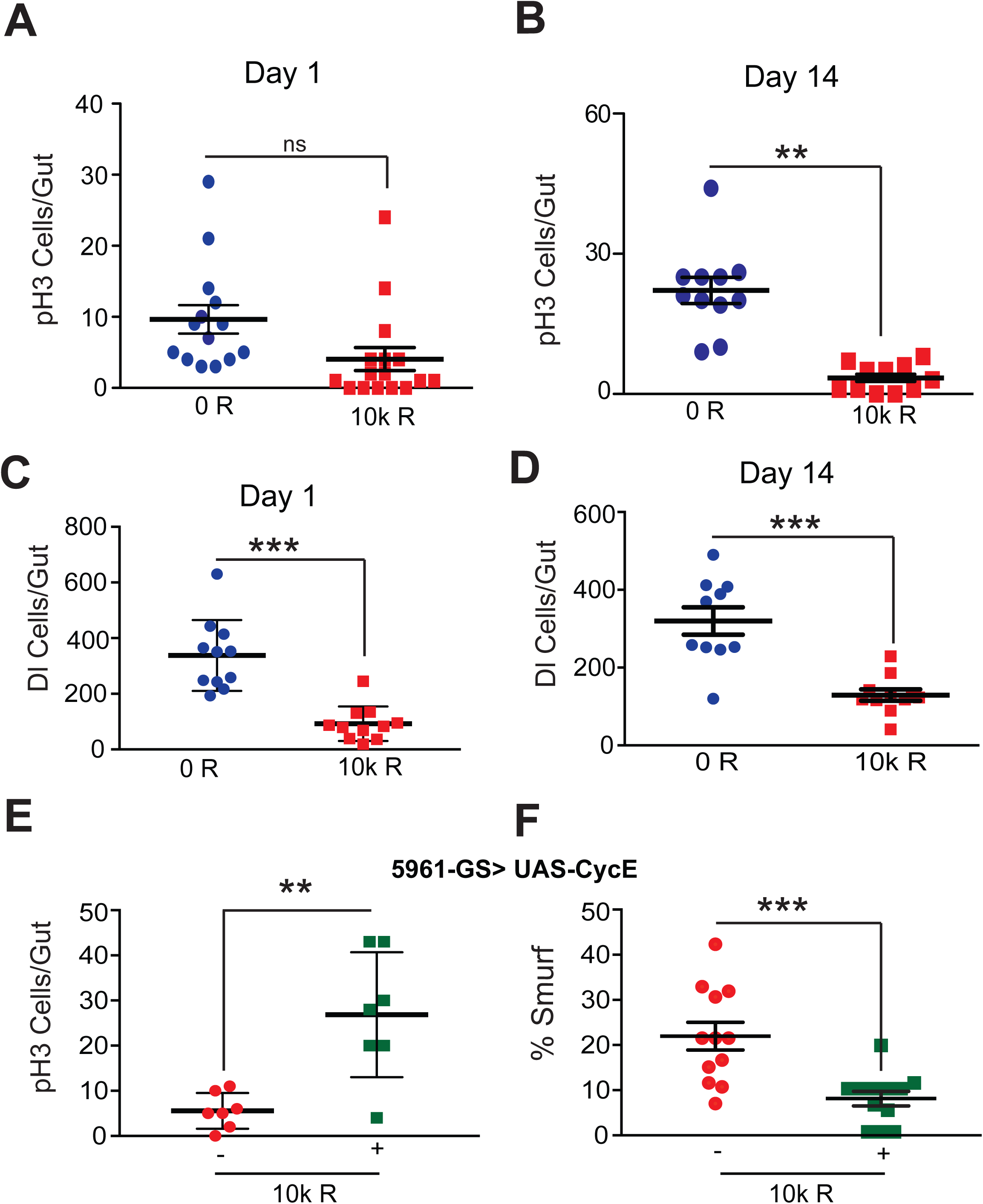
Radiation induced damage disrupts homeostatic proliferation by intestinal stem cells in the guts. **(A)** Intestinal stem cell proliferation was measured by counting the numbers of pH3-positive cells detected per gut of flies irradiated with 10k R day 1 and **(B)** day 14. Guts were dissected and pH3 staining was performed relevant days after irradiation with or without 10k R. The result is presented as mean ±SE of 10 guts per group. (** p < 0.01, * p < 0.05 by *t*-test). The effect of ISC proliferation on its ISCs was tested counting the numbers of β-Galactosidase positive cells Stem cell numbers were determined by immuno-staining using anti-β Gal antibody in dissected guts of lacZ enhancer trap line, P {PZ}Dl^05151^ (Dl-LacZ), in which the P-element is inserted at the 5′ un-translated region (UTR) of the *Delta* genomic locus on days 1 **(C)** and 14 **(D)** after irradiation. The result is presented as mean ±SE of 10 guts per group. (** p < 0.01, * p < 0.05 by *t*-test). **(E)** Number of pH3-positive cells detected per gut of *5961-GS>UAS-cycE* flies irradiated with 10k R. Guts were dissected and pH3 staining was performed 14 days after irradiation. Controls 10k R (-) maintained on 1.5 YE food without RU486, whereas cycE was overexpressed in ISCs in 10k R (+). The result is represented as mean ±SE of 10 guts per group. (** p < 0.01, * p < 0.05 by *t*-test). **(F)** Smurf assay for assessing gut permeability was performed with *5961-GS>UAS-cycE* flies on day 14 after irradiation. Control 10k R (-) maintained on 1.5 YE food without RU486, whereas cycE was overexpressed in ISCs in 10k R (+). Results plotted as mean proportion of Smurf to non-Smurf flies of 4 vials in each group with 25 flies in each vial. Error bars indicate SEM.

### Overexpression of *cyclin E* partially rescues radiation-induced intestinal permeability

Cyclin E/CDK2 plays a critical role in the G1 phase and the G1-S phase transition [80], and when overexpressed, overcomes cell cycle arrest [81]. We overexpressed *Cyclin E* (*CycE*) in ISCs (with the drug [RU486] inducible ISC-specific 5961 Gal4 UAS system) to test whether forced ISC proliferation rescues intestinal damage after irradiation. Flies where *CycE* was overexpressed (+), in an ISC-specific manner, had significantly increased ISC proliferation (as measured by pH3 staining in the intestine) when compared to the irradiated control without the RU486 (-) (**Fig. 3E).** We then enquired whether cycE over-expression in ISCs of irradiated flies also reduced intestinal permeability. We performed the Smurf assay in these flies and the results demonstrate about a 2-fold reduction in the percentage of flies with permeable guts, compared to the irradiated control (**Fig. 3F**). These results from the ectopic expression of *cyclin E* show that the effect of radiation damage to the intestine is reversible.

### Fly GWAS (Genome-Wide Association Study) for radiation-induced intestinal permeability

Genetic variations influence sensitivity to genotoxic stress, the detrimental effects of radiation treatment, and the prognosis of radiation therapy [83–92]. We hypothesized that *Drosophila* with naturally occurring genetic variations might reveal novel pathways that enhance stem cell proliferative repair and reduce radiation-induced intestinal permeability. To test this hypothesis, we leveraged the *Drosophila* Genetic Reference Panel that contains flies with fully sequenced genetic variations. We conducted an unbiased Genome Wide Association Study (GWAS screen) of approximately 150 fly strains from the *Drosophila* Genetic Reference Panel. Approximately 100 flies from each strain were irradiated with 10k R, and the percentage of Smurf flies was calculated 14 days after irradiation. The genetic markers with >25% minor allele frequency were used for screening [37]; lines were split into two groups, one for each allele at a given genetic locus. Linear regression modeling was used to determine the difference between phenotypes associated with each allele. The FDR for each trait was calculated by permutation of the phenotype data [44].

Results show that flies in different DGRP lines vary significantly in their susceptibility to radiation-induced intestinal damage. The DGRP lines varied in radiation-induced gut permeability, from a 14-fold increase in Smurf incidence to a 20-fold decrease in Smurf incidence (**Fig. 4**). Our GWAS analysis revealed several potential candidate genes (**Table 1**); however we setup a cutoff of false detection rate (FDR) of 27% or less to consider the genes for further validation [37]. We investigated the candidates listed in Table 1 for their ISC-specific influence on intestinal permeability after irradiation. To test this, we crossed fly lines expressing an RNAi against the candidate gene with virgin females with the drug-inducible (RU486) ISC-specific 5961 GS-driver. This allowed for temporal and spatial control over knockdown in the progeny. When the Smurf assay was performed in the 5-day-old adult female progeny (14 days after irradiation) among the candidates tested, we observed that knocking down *Msi* (*Musashi*) in ISCs most significantly increased gut permeability after irradiation (**Supplementary Fig. 3)**.

**Figure 4:**
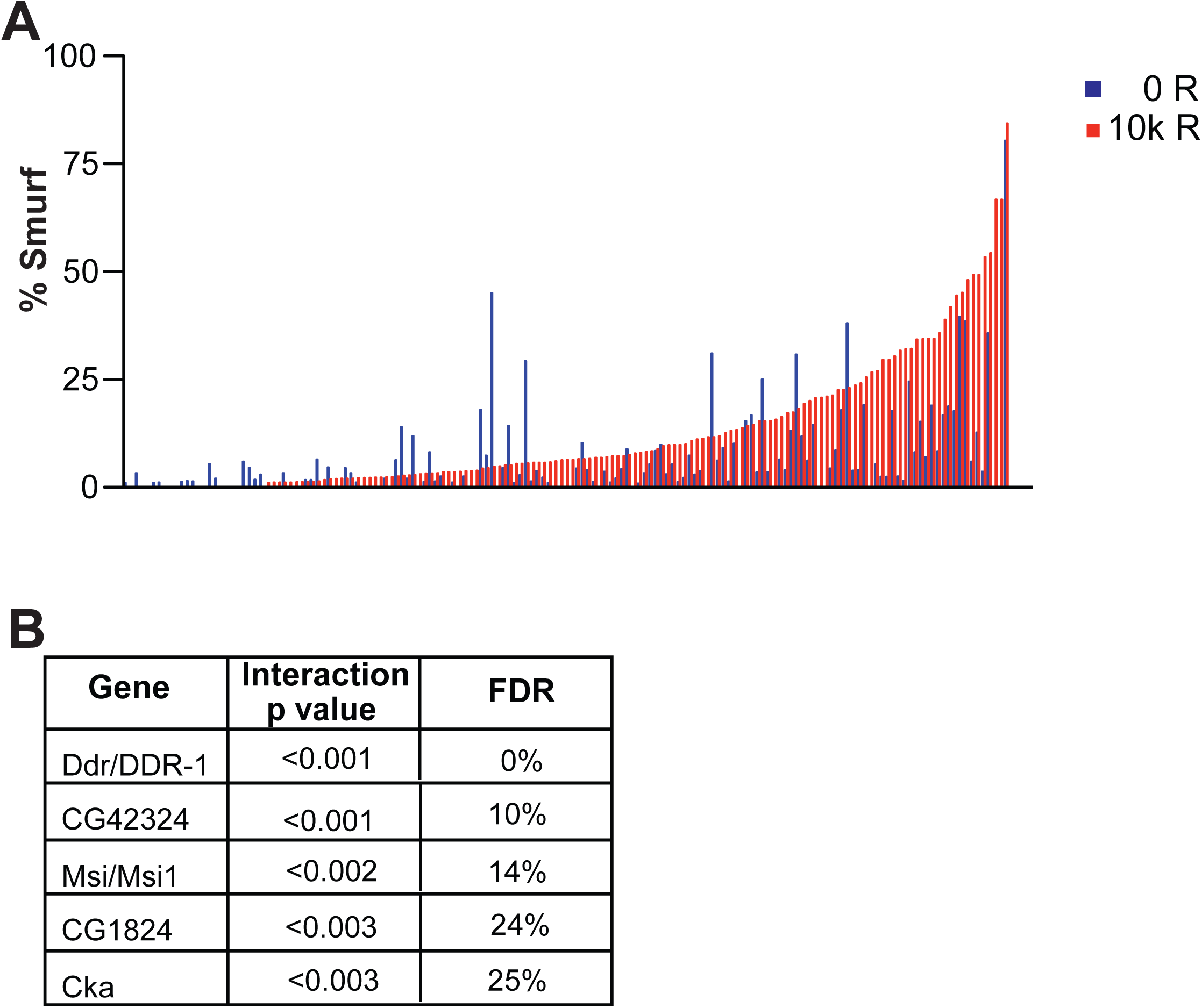
Phenotypic variation gut permeability across 180 DGRP lines caused by radiation exposure. **(A)** Lines are arranged in order of increasing phenotype of the irradiated lines, paired with their non-irradiated controls. Gut permeability, was determined by Smurf assay and results were plotted as mean proportion of ’Smurf’ to non-smurf flies in each group with at least 100 flies were tested per condition. (B) **Table 1**. List of candidate genes from a GWAS screen for gut permeability upon irradiation.

### *Musashi* regulates ISC function in response to radiation-induced damage

Musashi (Msi) belongs to a family of highly conserved RNA-binding translational repressors that are expressed in proliferative progenitor cells [93–95]. We hypothesized that ***Musashi (msi)*** modulates ISC proliferation in response to tissue damage from irradiation. To test this, we used the Gal4-UAS system to knock down msi expression ISCs [96]. The RNAi targeting *msi* was expressed using a drug-inducible (RU486) ISC-specific 5961 GS-driver in 5-day-old adult flies. Knocking down *msi* in ISCs (+), followed by irradiation, increased gut permeability by 2 times compared to irradiated control flies (without RU486) (-) exposed to 10k R **(Fig. 5A**). Conversely, when we overexpressed *msi* in ISCs (+) in irradiated flies, there was a significant reduction in gut permeability compared to irradiated control flies (without RU486) (-) **(Fig. 6A).** However, *msi* overexpression in enterocytes (EC) using 5966-Gal4 drivers did not rescue flies from gut permeability after irradiation, which supports an ISC-specific function for *msi* (**Supplementary Fig. S4A & B)**.

**Figure 5:**
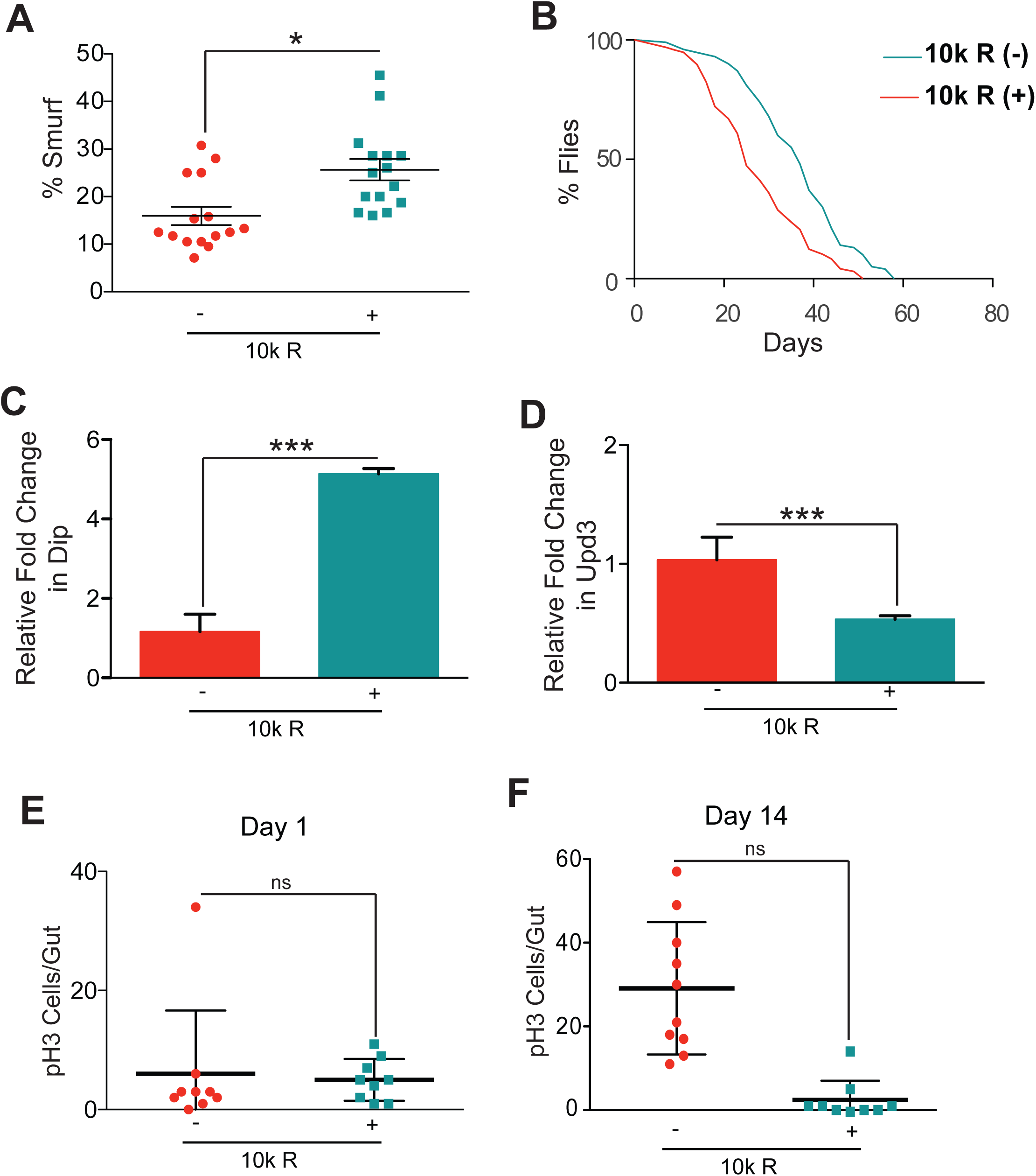
Reducing Msi expression in ISCs increases gut permeability and reduces survival by inhibiting stem cell proliferation. *5961-GS>UAS-Msi RNAi* flies were maintained in standard fly food with (+) to knockdown *msi* expression or without RU486 (-) in ISCs. Two days after adding RU486, flies were irradiated with 10k R to test the effect of its knockdown on intestinal homeostasis **(A)** Kaplan Meier survival analysis was performed for survival irradiation of 5 day old flies. At least 150 flies were used each group with control 10k R (-) maintained on 1.5 YE food without RU486, whereas Msi expression was knocked down in ISCs in 10k R (+). **(B)** Smurf assay for assessing gut permeability was performed day 14 after irradiation. Control 10k R (-) maintained on 1.5 YE food without RU486, whereas Msi was reduced in ISCs in 10k R (+). Results were plotted as mean change in the percentage of flies with permeable guts and the error bars indicate S.D. of 4 replicates. (** p < 0.01, * p < 0.05 by *t*-test). **(C)** Real time PCR to detect fold change in the expression of immune response genes *Diptericin* (*Dip*) in the dissected gut, 14 days after irradiation normalized to housekeeping gene RP49, control 10k R (-) maintained on 1.5 YE food without RU486, whereas Msi was knocked down in ISCs in 10k R (+). Results were plotted as mean fold change in expression and the error bars indicate S.D. of 4 replicates. (** p < 0.01, * p < 0.05 by *t*-test). **(D)** Real time PCR performed with RNA prepared from dissected guts, 24 hours after irradiation. The results are presented as mean fold change in the *Unpaired3* (*Upd3*) expression and normalized to housekeeping gene *RP49* and the error bars indicate S.D. of 4 replicates. (** p < 0.01, * p < 0.05 by *t*-test). Control 10k R (-) maintained on 1.5 YE food without RU486, whereas Msi was knocked down in 10k R (+). **(E)** The number of pH3-positive cells detected per gut of *5961-GS>UAS-Msi* flies, to knockdown Msi in ISCs (as indicated), on day 1 and **(F)** day 14 after irradiation in 10k R (+). Controls 10k R (-) maintained on 1.5 YE food without RU486. The result is represented as mean ±SE of 10 guts per group. (** p < 0.01, * p < 0.05 by *t*-test) of 3 independent experiments.

**Figure 6:**
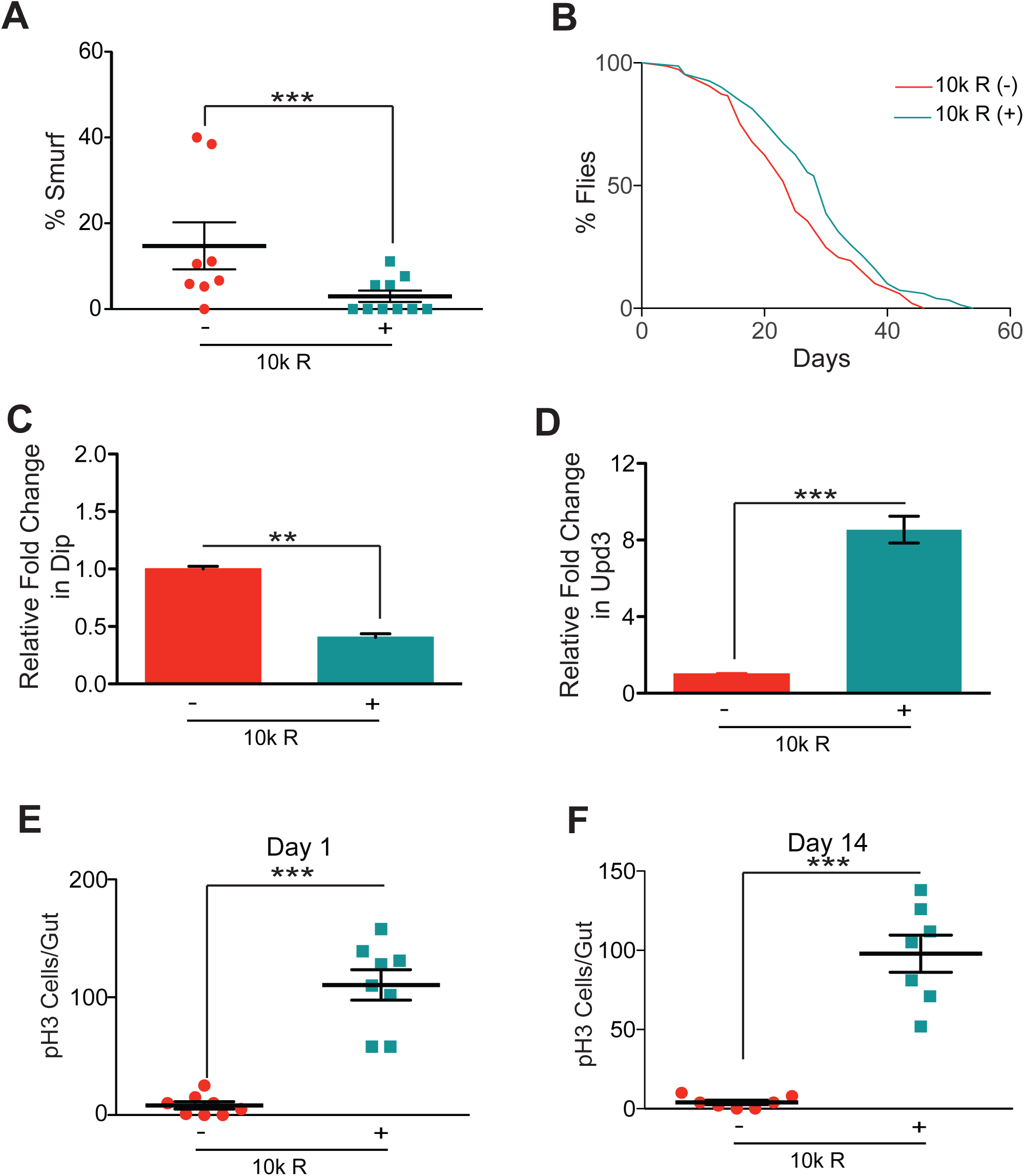
Increasing Msi in ISCs reduces gut permeability and increases survival by enhancing stem cell proliferation. *5961-GS>UAS-Msi* flies were maintained in standard fly food with (+) to over express *msi* expression or without RU486 (-) in ISCs. Two days after adding RU486, flies were irradiated with 10k R to test the effect of its knockdown on intestinal homeostasis **(A)** Kaplan Meier survival analysis was performed for survival irradiation of 5 day old flies. At least 150 flies were used each group with control 10k R (-) maintained on 1.5 YE food without RU486, whereas Msi expression was induced in ISCs in 10k R (+). **(B)** Smurf assay for assessing gut permeability was performed day 14 after irradiation. Control 10k R (-) maintained on 1.5 YE food without RU486, whereas Msi was over expressed in ISCs in 10k R (+). Results were plotted as mean change in the percentage of flies with permeable guts and the error bars indicate S.D. of 4 replicates. (** p < 0.01, * p < 0.05 by *t*-test). **(C)** Real time PCR to detect fold change in the expression of immune response genes *Diptericin* (*Dip*) in the dissected gut, 14 days after irradiation normalized to housekeeping gene RP49, control 10k R (-) maintained on 1.5 YE food without RU486, whereas Msi was over expressed in ISCs in 10k R (+). Results were plotted as mean fold change in expression and the error bars indicate S.D. of 4 replicates. (** p < 0.01, * p < 0.05 by *t*-test). **(D)** Real time PCR performed with RNA prepared from dissected guts, 24 hours after irradiation. The results are presented as mean fold change in the *Unpaired3* (*Upd3*) expression and normalized to housekeeping gene *RP49* and the error bars indicate S.D. of 4 replicates. (** p < 0.01, * p < 0.05 by *t*-test). Control 10k R (-) maintained on 1.5 YE food without RU486, whereas Msi was over expressed in 10k R (+). **(E)** The number of pH3-positive cells detected per gut of *5961-GS>UAS-Msi* flies, to induce Msi in ISCs (as indicated), on day 1 and **(F)** day 14 after irradiation in 10k R (+). Controls 10k R (-) maintained on 1.5 YE food without RU486. The result is represented as mean ±SE of 10 guts per group. (** p < 0.01, * p < 0.05 by *t*-test) of 3 independent experiments.

Since intestinal damage caused by radiation significantly reduced survival, we tested if *msi* expression improved survival in irradiated flies. Results indicate that *msi* knockdown in ISCs reduces survival upon irradiation **(Fig. 5B**), whereas *msi* overexpression results in a marginal but significant increase in survival **(Fig. 6B**). To further characterize msi’s impact on radiation sensitivity, we tested whether msi expression in ISC’s affected immune activation. In irradiated msi knockdown guts, qRT-PCR for *Dip,* a proxy for inflammation in the gut, showed a significant upregulation expression **(Fig. 5C**). Conversely, dissected intestines with ISC-specific msi overexpression, showed a a significant reduction in *Dip* expression **(Fig. 6C**). Interestingly, real-time PCR in the same dissected guts indicated a reduced expression of Upd3 upon *msi* knockdown **(Fig. 5D**), whereas *msi* overexpression correlated with a significant increase in Upd3 expression in the gut 24 hours after irradiation **(Fig. 6D**).

Because *msi* is known to regulate cell fate and stemness [97], and irradiation significantly reduces ISC proliferation, we investigated the effect of *msi* on ISC proliferation in response to radiation. Results indicate that when *msi* was knocked down in ISCs, ISC proliferation, as measured by immunofluorescence for **phospho-Histone 3 (pH3)** in dissected guts, was comparable to the control 1 and 14 days after irradiation (-) **(Fig. 5E & F**). However, pH3 immunofluorescence staining demonstrated that *msi* overexpression significantly increased ISC proliferation by 15 times **(Fig. 6E**). Also, flies overexpressing *msi* showed a sustained elevation in the number of pH3 positive cells in the gut, even 14 days after irradiation, albeit half of day one levels **(Fig. 6F**). Thus the survival of flies upon manipulating *msi* levels was correlated with its ability to influence ISC proliferation.

## Discussion

Understanding the mechanisms involved in tissues homeostasis and repair in response to genotoxic stress is critical for understanding how organisms deal with age-related accumulation of genotoxic stress. This knowledge may also help in developing therapeutics against the side effects of chemotherapeutic agents and However, the lack of an effective *in vivo* model has hampered progress. We have developed adult *Drosophila melanogaster* as a model to study how different cells interact to mount a response to genotoxic stress to maintain tissue homeostasis and repair. We leveraged the conservation of the fly intestine to characterize the effect of ionizing radiation on tissue homeostasis and screened naturally occurring genetic variants to identify alleles germane to intestinal permeability. In addition, we utilized naturally occurring genetic variations in flies to identify novel regulators of tissue damage caused by DNA damage. Our GWAS analysis in these strains identified musashi (*msi*) as a potential candidate. Further results showed that the levels of *msi* in ISCs correlated with ISC proliferation and ectopic expression of msi in ISC not only reduced intestinal permeability but also increased survival in response to irradiation.

Earlier studies have shown that exposure to radiation in adult female flies affected fecundity and chromosomal aberrations in the progeny [98]. However, little was known regarding the long-term effect of ionizing radiation on survival of adult flies. We began by investigating the dosage of radiation that reduces survival. Our results demonstrate that flies are quite resistant to tissue damage caused by ionizing radiation, which is in agreement to previously published literature [99, 100]. We find that when flies are exposed to either a staggered or singular dose (4 doses of 2500 R every other day) of 10K R, gut permeability is enhanced and survival is reduced. Exposing flies to staggered doses of radiation, may be more representative of patients undergoing radiation therapy. The effect of radiation on survival was independent of sex as results were consistent between male and female flies. Even though 10k R is higher than doses tolerated by mammals, it is lower compared to past studies where flies were irradiated with ionizing radiation [101]. This as discussed earlier is on account of flies possessing mechanisms that protects its dividing cells from genotoxic stress that are not analogous in human cells [102].

Since we exposed whole flies to radiation we expected a strong physiological readout that might explain shortened survival of irradiated flies. We observed a consistent increase in the phosphorylation of γ-H2Av, the fly orthologue of H2AX [66]. We also observed elevated intestinal permeability and shorter small intestines. As increased gut permeability has previously been associated with reduced survival [60, 103] due to increased local and systemic inflammation we performed semi-quantitative Smurf assay which demonstrated indeed irradiated flies have highly permeable intestines. In addition, we also observed elevated inflammation in the intestine quite early after irradiation, followed by increased systemic inflammation that temporally correlated with increased intestinal permeability (by day 14 after irradiation). We speculate that increased intestinal permeability and disruption of barrier function leading to exposure to commensal micro flora might explain highly up regulated systemic inflammation as a potential cause for lethality a few weeks after irradiation. In fact our anecdotal observation indicated that Smurf flies were more likely to dye.

In our model, there are two lines of evidence that suggest increased JNK signaling in response to radiation: One, there is an increase in expression of the IL-6-like cytokine, Upd3, in response to radiation [71, 104] Two, there is a radiation-induced expression of *Hid, reaper* and *puckered* in the intestines, which presumably leads to apoptosis in the gut [105]. In humans, the detrimental responses to radiation treatment vary greatly [107, 108] and survival, health, and gut homeostasis may at least in part be regulated by genetic factors [85, 88, 89]. Interestingly, leaky gut syndrome is a hallmark of radiation enteritis in human patients undergoing radiation therapy [107, 108]. Thus, we reasoned if fully sequenced natural variations amongst various DGRP strains might be used for discovering novel genes that may restore intestinal stem cell function in irradiated flies. We chose the Smurf assay as the readout for our screen because it is a quantitative measure of intestinal health. Previously, a case-control analysis in DGRP lines for radiation resistance identified several candidate genes with human orthologue [109], however they only evaluated genes involved in survival following acute exposure to radiation and did not account for tissue damage and repair several weeks after exposure. In addition, our linear regression analysis is also more robust compared to case-control analysis in limiting confounding [110].

Interestingly, we observed a significant decline in proliferating ISCs, which reduced ISCs numbers. So we reasoned that the dual effect of radiation on increased apoptosis in the intestine and reduction in reparative proliferation might be responsible for increased intestinal permeability in irradiated flies. When then asked if forcing the restoration of ISC populations might have a protective effect in irradiated flies. Previous studies have shown that *cyclin E* is involved with cells’ re-entry into the cell cycle. In flies, *Cyclin E* alone is capable of activating re-entry into S-phase and promotes ISC proliferation [106]. In addition, overexpression of *cyclin E* promotes proliferation in cells while Ras activation and mitochondrial dysfunction mediates reactive oxygen species production, leading to activation of p53 and cell cycle arrest [81]. Our results confirmed that over-expression of cycE in ISCs not only restored intestinal barrier function, it also increased survival and that observed effect of radiation induced tissue damage maybe reversible.

*Musashi* (*msi*) *is* a highly conserved RNA binding protein regulator of post-transcriptional processing of target genes [38], as well as a known stem cell marker [97]. It was first identified as a regulator of asymmetric division sensory organ precursor cells in *Drosophila* [40]. We found that modulating *msi* in ISCs affected ISC proliferation, which is consistent with the human orthologue, *msi1* that is strongly expressed in the intestinal crypts, especially during embryonic development and regeneration [94]. Interestingly, *msi* overexpression did not significantly impact survival in non-irradiated flies. The stem cell specific role of msi was further confirmed since its ectopic expression in enterocytes had no effect on intestinal permeability. Interestingly, *msi1* knockdown in U-251 (human glioblastoma cell line) resulted in higher instances of double-stranded breaks [111], suggesting its role in DNA repair. We were surprised to observe that msi over expression in ISCs resulted in increased expression of Upd3 expression in dissected guts from irradiated flies. Since Upd3 is expressed primarily in ECs, we suspect this maybe sign of healthier intestine upon msi over-expression 2 days before irradiation. However it may also suggest that a more intriguing possibility that msi maybe involved in a potential cross talk between ISCs and ECs, which remains to be tested.

Another study in mice demonstrated that *msi1* and *msi2* could regulate stem cell activation and self-renewal of crypt base columnar cells upon tissue damage, thus indicating a conserved effect of *msi* on ISC function [112] none the less our findings in conjunction to these reports points to critical role of musashi in regenerative medicine.

## Figure legends

**Supplementary Figure 1: Radiation induced damage results in significant reduction in survival and increased intestinal permeability in dose dependent manner: (A)** Kaplan Meier survival analysis for survival irradiation of 5 day old flies. At least 150 flies were used for each dose. **(B)** Representative image of the dissected intestine of adult female fly non-irradiated (0 R) control flies and irradiated (10k R) female flies 14 days after irradiation. **(C)** Kaplan Meier survival analysis for survival irradiation of 5 day old flies male *w*^*1118*^ flies, showing the effect of radiation on survival is independent of sex. At least 150 flies were used for control (0R) and flies irradiated with 10k Rads (10kR). **(D)** The Smurf assay was performed to investigate the effect of radiation on gut permeability in *w*^*1118*^ female flies after irradiation with or without 500 rads, 1000 rads, 2000 rads and 5000 rads (0.5k R, 1k R, 2k R, 5k R, 0 R as indicated) at 14 days after irradiation. The results were plotted as mean percentage of ‘Smurf’ to non-smurf flies. Error bars indicate S.D. of 4 replicates. (** p < 0.01, * p < 0.05 by *t*-test). (E) The Smurf assay was performed to investigate the effect of radiation on gut permeability. The adult female *w*^*1118*^ 5 days old adult flies were irradiated with 10k rads (10k R) or flies received ‘staggered’ dosage of 2000 rads every other day for 5 days (10k R (S)) and compared to non-irradiated control (0 R). Smurf assay was performed 14 days after first dose and results were plotted as mean percentage of ‘Smurf’ to non-smurf flies. Error bars indicate S.D. of 4 replicates. (** p < 0.01, * p < 0.05 by *t*-test). (F) The Smurf assay was performed to investigate the effect of radiation on gut permeability in either male or female flies 14 after irradiation (10k R) and percentage of Smurf flies were compared to non-irradiated control (0 R). Smurf assay was performed 14 days after first dose and results were plotted as mean percentage of ’Smurf’ to non-smurf flies. Error bars indicate S.D. of 4 replicates. (** p < 0.01, * p < 0.05 by *t*-test).

**Supplementary Figure 2: Exposure to radiation causes DNA damage and cell death in dose dependent manner. (A)** Representative image of adult *Drosophila* gut dissected 30 minutes after irradiation with or without 0 R, 1k R, 2kR or 5k R (irradiated with 1000, 2000 and 5000 rads respectively) and stained with γ-H2Av antibody (Red) and co-stained with DAPI (blue) **(B)** Mean fluorescence intensity of GFP expression from images of the dissected gut from recombinant line, *Upd-3-Gal4 UAS-GFP*, 3 days after irradiation with or without 10k R. Error bars indicate S.D. of 5 replicates. (** p < 0.01, * p < 0.05 by *t*-test). **(C)** Real time PCR performed with RNA prepared from dissected guts, 24 hours after irradiation (10k R) or non-irradiated control (0 R). The results are presented as mean fold change in the *Unpaired3* (*Upd3*) expression and normalized to housekeeping gene *RP49* and the error bars indicate S.D. of 4 replicates. (** p < 0.01, * p < 0.05 by *t*-test). **(D)** Ethidium bromide-Acridine Orange staining was performed in guts of *w*^*1118*^ female flies after irradiation with or without 1000 rads, 2000 rads and 5000 rads (1k R, 2k R, 5k R, or control (0 R) as indicated at 48 hours after irradiation. The results are plotted as the mean number of apoptotic cells per gut and presented and mean apoptotic cells and error bars indicate S.D. of 2 independent experiments with at least 5 guts each.

**Supplementary Figure 3: Secondary screen for intestinal permeability for candidate genes from DGRP lines smurf assay.** The expression of candidate genes was knocked down in ISCs using *5961-GS driver* (as indicated). *5961-GS>UAS-candidate gene* flies were maintained in standard fly food with (+) to knockdown its expression or without RU486 (-). Two days after adding RU486, flies were irradiated with 10k R to test the effect of its knockdown on intestinal permeability. Results were plotted as mean change in the percentage of flies with permeable guts and the error bars indicate S.D. of 4 replicates. (** p < 0.01, * p < 0.05 by *t*-test).

## Acknowledgments

We thank the Bloomington Stock Center and Vienna *Drosophila* RNAi Center for providing the fly strains. We also thank members of Kapahi and Jasper labs for discussions and suggestions. We would also like to thank Geoffrey Meyerhof for his help editing the manuscript. This work was funded by grants from the American Federation of Aging Research and Hillblom foundations (P.K., S.D.K.), and the NIH (R01AG038688 & RO1AG045835) (P.K.).

## References

1. Barnes DE, Lindahl T. Repair and genetic consequences of endogenous DNA base damage in mammalian cells. Annu Rev Genet. 2004;38:445–76. doi: 10.1146/annurev.genet.38.072902.092448. PubMed PMID: 15568983.

2. Azzam EI, Jay-Gerin JP, Pain D. Ionizing radiation-induced metabolic oxidative stress and prolonged cell injury. Cancer letters. 2012;327(1-2):48–60. doi: 10.1016/j.canlet.2011.12.012. PubMed PMID: 22182453; PubMed Central PMCID: PMCPMC3980444.

3. Li H, Mitchell JR, Hasty P. DNA double-strand breaks: a potential causative factor for mammalian aging? Mechanisms of ageing and development. 2008;129(7-8):416–24. doi: 10.1016/j.mad.2008.02.002. PubMed PMID: 18346777; PubMed Central PMCID: PMCPMC2517577.

4. White RR, Vijg J. Do DNA Double-Strand Breaks Drive Aging? Molecular cell. 2016;63(5):729–38. doi: 10.1016/j.molcel.2016.08.004. PubMed PMID: 27588601; PubMed Central PMCID: PMCPMC5012315.

5. Zhang D, Wang HB, Brinkman KL, Han SX, Xu B. Strategies for targeting the DNA damage response for cancer therapeutics. Chin J Cancer. 2012;31(8):359–63. doi: 10.5732/cjc.012.10087. PubMed PMID: 22704491; PubMed Central PMCID: PMCPMC3777509.

6. Ljungman M. Targeting the DNA damage response in cancer. Chem Rev. 2009;109(7):2929–50. doi: 10.1021/cr900047g. PubMed PMID: 19545147.

7. Powell SN, Bindra RS. Targeting the DNA damage response for cancer therapy. DNA repair. 2009;8(9):1153–65. doi: 10.1016/j.dnarep.2009.04.011. PubMed PMID: 19501553.

8. O’Connor MJ. Targeting the DNA Damage Response in Cancer. Molecular cell. 2015;60(4):547–60. doi: 10.1016/j.molcel.2015.10.040. PubMed PMID: 26590714.

9. Castedo M, Perfettini JL, Roumier T, Andreau K, Medema R, Kroemer G. Cell death by mitotic catastrophe: a molecular definition. Oncogene. 2004;23(16):2825–37. Epub 2004/04/13. doi: 10.1038/sj.onc.1207528. PubMed PMID: 15077146.

10. Vaupel P. Tumor microenvironmental physiology and its implications for radiation oncology. Seminars in radiation oncology. 2004;14(3):198–206. Epub 2004/07/16. doi: 10.1016/j.semradonc.2004.04.008. PubMed PMID: 15254862.

11. Gomez-Millan J, Katz IS, Farias Vde A, Linares-Fernandez JL, Lopez-Penalver J, Ortiz-Ferron G, et al. The importance of bystander effects in radiation therapy in melanoma skin-cancer cells and umbilical-cord stromal stem cells. Radiotherapy and oncology : journal of the European Society for Therapeutic Radiology and Oncology. 2012;102(3):450–8. Epub 2011/12/16. doi: 10.1016/j.radonc.2011.11.002. PubMed PMID: 22169765.

12. Joye I, Haustermans K. Early and late toxicity of radiotherapy for rectal cancer. Recent results in cancer research Fortschritte der Krebsforschung Progres dans les recherches sur le cancer. 2014;203:189–201. Epub 2014/08/12. doi: 10.1007/978-3-319-08060-4_13. PubMed PMID: 25103006.

13. Zhu Y, Zhou J, Tao G. Molecular aspects of chronic radiation enteritis. Clinical and investigative medicine Medecine clinique et experimentale. 2011;34(3):E119–24. Epub 2011/06/03. PubMed PMID: 21631987.

14. Accordino MK, Neugut AI, Hershman DL. Cardiac effects of anticancer therapy in the elderly. Journal of clinical oncology : official journal of the American Society of Clinical Oncology. 2014;32(24):2654–61. Epub 2014/07/30. doi: 10.1200/JCO.2013.55.0459. PubMed PMID: 25071122.

15. Bourgier C, Lacombe J, Solassol J, Mange A, Pelegrin A, Ozsahin M, et al. Late side-effects after curative intent radiotherapy: Identification of hypersensitive patients for personalized strategy. Critical reviews in oncology/hematology. 2014. Epub 2014/12/17. doi: 10.1016/j.critrevonc.2014.11.004. PubMed PMID: 25497158.

16. Kelley MR, Logsdon D, Fishel ML. Targeting DNA repair pathways for cancer treatment: what’s new? Future Oncol. 2014;10(7):1215–37. doi: 10.2217/fon.14.60. PubMed PMID: 24947262; PubMed Central PMCID: PMCPMC4125008.

17. Howard SM, Yanez DA, Stark JM. DNA damage response factors from diverse pathways, including DNA crosslink repair, mediate alternative end joining. PLoS genetics. 2015;11(1):e1004943. doi: 10.1371/journal.pgen.1004943. PubMed PMID: 25629353; PubMed Central PMCID: PMCPMC4309583.

18. Liu C, Niu Y, Zhou X, Xu X, Yang Y, Zhang Y, et al. Cell cycle control, DNA damage repair, and apoptosis-related pathways control pre-ameloblasts differentiation during tooth development. BMC Genomics. 2015;16:592. doi: 10.1186/s12864-015-1783-y. PubMed PMID: 26265206; PubMed Central PMCID: PMCPMC4534026.

19. Corcoran NM, Clarkson MJ, Stuchbery R, Hovens CM. Molecular Pathways: Targeting DNA Repair Pathway Defects Enriched in Metastasis. Clin Cancer Res. 2016;22(13):3132–7. doi: 10.1158/1078-0432.CCR-15-1050. PubMed PMID: 27169997.

20. d’Adda di Fagagna F. Living on a break: cellular senescence as a DNA-damage response. Nature reviews Cancer. 2008;8(7):512–22. doi: 10.1038/nrc2440. PubMed PMID: 18574463.

21. Childs BG, Baker DJ, Kirkland JL, Campisi J, van Deursen JM. Senescence and apoptosis: dueling or complementary cell fates? EMBO Rep. 2014;15(11):1139–53. doi: 10.15252/embr.201439245. PubMed PMID: 25312810; PubMed Central PMCID: PMCPMC4253488.

22. Stacey R, Green JT. Radiation-induced small bowel disease: latest developments and clinical guidance. Ther Adv Chronic Dis. 2014;5(1):15–29. doi: 10.1177/2040622313510730. PubMed PMID: 24381725; PubMed Central PMCID: PMCPMC3871275.

23. Shadad AK, Sullivan FJ, Martin JD, Egan LJ. Gastrointestinal radiation injury: prevention and treatment. World journal of gastroenterology : WJG. 2013;19(2):199–208. doi: 10.3748/wjg.v19.i2.199. PubMed PMID: 23345942; PubMed Central PMCID: PMCPMC3547575.

24. Leulier F, Royet J. Maintaining immune homeostasis in fly gut. Nature immunology. 2009;10(9):936–8. Epub 2009/08/21. doi: 10.1038/ni0909-936. PubMed PMID: 19692992.

25. Ochoa-Reparaz J, Mielcarz DW, Begum-Haque S, Kasper LH. Gut, bugs, and brain: role of commensal bacteria in the control of central nervous system disease. Annals of neurology. 2011;69(2):240–7. Epub 2011/03/10. doi: 10.1002/ana.22344. PubMed PMID: 21387369.

26. Micchelli CA, Perrimon N. Evidence that stem cells reside in the adult Drosophila midgut epithelium. Nature. 2006;439(7075):475–9. Epub 2005/12/13. doi: 10.1038/nature04371. PubMed PMID: 16340959.

27. Lane SW, Williams DA, Watt FM. Modulating the stem cell niche for tissue regeneration. Nature biotechnology. 2014;32(8):795–803. Epub 2014/08/06. doi: 10.1038/nbt.2978. PubMed PMID: 25093887.

28. Ayyaz A, Jasper H. Intestinal inflammation and stem cell homeostasis in aging Drosophila melanogaster. Frontiers in cellular and infection microbiology. 2013;3:98. Epub 2014/01/01. doi: 10.3389/fcimb.2013.00098. PubMed PMID: 24380076; PubMed Central PMCID: PMC3863754.

29. Chatterjee M, Ip YT. Pathogenic stimulation of intestinal stem cell response in Drosophila. Journal of cellular physiology. 2009;220(3):664–71. Epub 2009/05/20. doi: 10.1002/jcp.21808. PubMed PMID: 19452446; PubMed Central PMCID: PMC4003914.

30. Zhang L, Sun W, Wang J, Zhang M, Yang S, Tian Y, et al. Mitigation effect of an FGF-2 peptide on acute gastrointestinal syndrome after high-dose ionizing radiation. International journal of radiation oncology, biology, physics. 2010;77(1):261–8. Epub 2010/04/17. doi: 10.1016/j.ijrobp.2009.11.026. PubMed PMID: 20394858; PubMed Central PMCID: PMC2883168.

31. Qiu W, Carson-Walter EB, Liu H, Epperly M, Greenberger JS, Zambetti GP, et al. PUMA regulates intestinal progenitor cell radiosensitivity and gastrointestinal syndrome. Cell stem cell. 2008;2(6):576–83. Epub 2008/06/05. doi: 10.1016/j.stem.2008.03.009. PubMed PMID: 18522850; PubMed Central PMCID: PMC2892934.

32. Koukourakis MI. Radiation damage and radioprotectants: new concepts in the era of molecular medicine. The British journal of radiology. 2012;85(1012):313–30. Epub 2012/02/02. doi: 10.1259/bjr/16386034. PubMed PMID: 22294702; PubMed Central PMCID: PMC3486665.

33. Poglio S, Galvani S, Bour S, Andre M, Prunet-Marcassus B, Penicaud L, et al. Adipose tissue sensitivity to radiation exposure. The American journal of pathology. 2009;174(1):44–53. Epub 2008/12/20. doi: 10.2353/ajpath.2009.080505. PubMed PMID: 19095959; PubMed Central PMCID: PMC2631317.

34. Wang L, Jones DL. The effects of aging on stem cell behavior in Drosophila. Experimental gerontology. 2011;46(5):340–4. doi: 10.1016/j.exger.2010.10.005. PubMed PMID: 20971182; PubMed Central PMCID: PMCPMC3079770.

35. Hong AW, Meng Z, Guan KL. The Hippo pathway in intestinal regeneration and disease. Nature reviews Gastroenterology & hepatology. 2016;13(6):324–37. doi: 10.1038/nrgastro.2016.59. PubMed PMID: 27147489; PubMed Central PMCID: PMCPMC5642988.

36. Kux K, Pitsouli C. Tissue communication in regenerative inflammatory signaling: lessons from the fly gut. Frontiers in cellular and infection microbiology. 2014;4:49. doi: 10.3389/fcimb.2014.00049. PubMed PMID: 24795868; PubMed Central PMCID: PMCPMC4006025.

37. Mackay TF, Richards S, Stone EA, Barbadilla A, Ayroles JF, Zhu D, et al. The Drosophila melanogaster Genetic Reference Panel. Nature. 2012;482(7384):173–8. doi: 10.1038/nature10811. PubMed PMID: 22318601; PubMed Central PMCID: PMC3683990.

38. Zearfoss NR, Deveau LM, Clingman CC, Schmidt E, Johnson ES, Massi F, et al. A conserved three-nucleotide core motif defines Musashi RNA binding specificity. The Journal of biological chemistry. 2014;289(51):35530–41. doi: 10.1074/jbc.M114.597112. PubMed PMID: 25368328; PubMed Central PMCID: PMCPMC4271237.

39. Siddall NA, McLaughlin EA, Marriner NL, Hime GR. The RNA-binding protein Musashi is required intrinsically to maintain stem cell identity. Proceedings of the National Academy of Sciences of the United States of America. 2006;103(22):8402–7. doi: 10.1073/pnas.0600906103. PubMed PMID: 16717192; PubMed Central PMCID: PMCPMC1570104.

40. Nakamura M, Okano H, Blendy JA, Montell C. Musashi, a neural RNA-binding protein required for Drosophila adult external sensory organ development. Neuron. 1994;13(1):67–81. PubMed PMID: 8043282.

41. Zid BM, Rogers AN, Katewa SD, Vargas MA, Kolipinski MC, Lu TA, et al. 4E-BP extends lifespan upon dietary restriction by enhancing mitochondrial activity in Drosophila. Cell. 2009;139(1):149–60. doi: 10.1016/j.cell.2009.07.034. PubMed PMID: 19804760; PubMed Central PMCID: PMCPMC2759400.

42. Kapahi P, Zid BM, Harper T, Koslover D, Sapin V, Benzer S. Regulation of lifespan in Drosophila by modulation of genes in the TOR signaling pathway. Curr Biol. 2004;14(10):885–90. doi: 10.1016/j.cub.2004.03.059. PubMed PMID: 15186745; PubMed Central PMCID: PMCPMC2754830.

43. Katewa SD, Demontis F, Kolipinski M, Hubbard A, Gill MS, Perrimon N, et al. Intramyocellular fatty-acid metabolism plays a critical role in mediating responses to dietary restriction in Drosophila melanogaster. Cell metabolism. 2012;16(1):97–103. doi: 10.1016/j.cmet.2012.06.005. PubMed PMID: 22768842; PubMed Central PMCID: PMC3400463.

44. Palma-Guerrero J, Hall CR, Kowbel D, Welch J, Taylor JW, Brem RB, et al. Genome wide association identifies novel loci involved in fungal communication. PLoS genetics. 2013;9(8):e1003669. doi: 10.1371/journal.pgen.1003669. PubMed PMID: 23935534; PubMed Central PMCID: PMC3731230.

45. !!! INVALID CITATION !!! (67).

46. Abrahamson S, Friedman LD. X-Ray Induced Mutations in Spermatogonial Cells of Drosophila and Their Dose-Frequency Relationship. Genetics. 1964;49:357–61. Epub 1964/02/01. PubMed PMID: 14124949; PubMed Central PMCID: PMC1210575.

47. Eeken JC, de Jong AW, Loos M, Vreeken C, Romeyn R, Pastink A, et al. The nature of X-ray-induced mutations in mature sperm and spermatogonial cells of Drosophila melanogaster. Mutation research. 1994;307(1):201–12. Epub 1994/05/01. PubMed PMID: 7513798.

48. Bownes M, Sunnell LA. Developmental effects of X-irradiation of early Drosophila embryos. Journal of embryology and experimental morphology. 1977;39:253–9. Epub 1977/06/01. PubMed PMID: 407324.

49. Parashar V, Frankel S, Lurie AG, Rogina B. The effects of age on radiation resistance and oxidative stress in adult Drosophila melanogaster. Radiat Res. 2008;169(6):707–11. doi: 10.1667/RR1225.1. PubMed PMID: 18494545.

50. Noal S, Levy C, Hardouin A, Rieux C, Heutte N, Segura C, et al. One-year longitudinal study of fatigue, cognitive functions, and quality of life after adjuvant radiotherapy for breast cancer. International journal of radiation oncology, biology, physics. 2011;81(3):795–803. Epub 2010/10/05. doi: 10.1016/j.ijrobp.2010.06.037. PubMed PMID: 20888704.

51. Gulliford SL, Miah AB, Brennan S, McQuaid D, Clark CH, Partridge M, et al. Dosimetric explanations of fatigue in head and neck radiotherapy: an analysis from the PARSPORT Phase III trial. Radiotherapy and oncology : journal of the European Society for Therapeutic Radiology and Oncology. 2012;104(2):205–12. Epub 2012/08/14. doi: 10.1016/j.radonc.2012.07.005. PubMed PMID: 22883107.

52. Courtier N, Gambling T, Enright S, Barrett-Lee P, Abraham J, Mason MD. A prognostic tool to predict fatigue in women with early-stage breast cancer undergoing radiotherapy. Breast. 2013;22(4):504–9. Epub 2012/10/30. doi: 10.1016/j.breast.2012.10.002. PubMed PMID: 23103133.

53. Geinitz H, Zimmermann FB, Stoll P, Thamm R, Kaffenberger W, Ansorg K, et al. Fatigue, serum cytokine levels, and blood cell counts during radiotherapy of patients with breast cancer. International journal of radiation oncology, biology, physics. 2001;51(3):691–8. Epub 2001/10/13. PubMed PMID: 11597810.

54. Smets EM, Visser MR, Willems-Groot AF, Garssen B, Oldenburger F, van Tienhoven G, et al. Fatigue and radiotherapy: (A) experience in patients undergoing treatment. British journal of cancer. 1998;78(7):899–906. Epub 1998/10/09. PubMed PMID: 9764581; PubMed Central PMCID: PMC2063131.

55. Faithfull S. Fatigue in patients receiving radiotherapy. Professional nurse. 1998;13(7):459–61. Epub 1998/07/08. PubMed PMID: 9653282.

56. Renner M, Feng R, Springer D, Chen MK, Ntamack A, Espina A, et al. A murine model of peripheral irradiation-induced fatigue. Behav Brain Res. 2016;307:218–26. doi: 10.1016/j.bbr.2016.03.035. PubMed PMID: 27012391; PubMed Central PMCID: PMCPMC4853268.

57. Wolff BS, Renner MA, Springer DA, Saligan LN. A Mouse Model of Fatigue Induced by Peripheral Irradiation. J Vis Exp. 2017;(121). doi: 10.3791/55145. PubMed PMID: 28362365.

58. Pfeiffenberger C, Lear BC, Keegan KP, Allada R. Locomotor activity level monitoring using the Drosophila Activity Monitoring (DAM) System. Cold Spring Harb Protoc. 2010;2010(11):pdb prot5518. doi: 10.1101/pdb.prot5518. PubMed PMID: 21041391.

59. Pereira MT, Malik M, Nostro JA, Mahler GJ, Musselman LP. Effect of dietary additives on intestinal permeability in both Drosophila and a human cell co-culture. Disease models & mechanisms. 2018;11(12). doi: 10.1242/dmm.034520. PubMed PMID: 30504122.

60. Rera M, Clark RI, Walker DW. Intestinal barrier dysfunction links metabolic and inflammatory markers of aging to death in Drosophila. Proceedings of the National Academy of Sciences of the United States of America. 2012;109(52):21528–33. doi: 10.1073/pnas.1215849110. PubMed PMID: 23236133; PubMed Central PMCID: PMCPMC3535647.

61. Tzou P, Ohresser S, Ferrandon D, Capovilla M, Reichhart JM, Lemaitre B, et al. Tissue-specific inducible expression of antimicrobial peptide genes in Drosophila surface epithelia. Immunity. 2000;13(5):737–48. PubMed PMID: 11114385.

62. Ferrandon D, Jung AC, Criqui M, Lemaitre B, Uttenweiler-Joseph S, Michaut L, et al. A drosomycin-GFP reporter transgene reveals a local immune response in Drosophila that is not dependent on the Toll pathway. The EMBO journal. 1998;17(5):1217–27. Epub 1998/04/18. doi: 10.1093/emboj/17.5.1217. PubMed PMID: 9482719; PubMed Central PMCID: PMC1170470.

63. Meng X, Khanuja BS, Ip YT. Toll receptor-mediated Drosophila immune response requires Dif, an NF-kappaB factor. Genes Dev. 1999;13(7):792–7. PubMed PMID: 10197979; PubMed Central PMCID: PMCPMC316597.

64. Hetru C, Hoffmann JA. NF-kappaB in the immune response of Drosophila. Cold Spring Harb Perspect Biol. 2009;1(6):a000232. doi: 10.1101/cshperspect.a000232. PubMed PMID: 20457557; PubMed Central PMCID: PMCPMC2882123.

65. Tang H, Kambris Z, Lemaitre B, Hashimoto C. A serpin that regulates immune melanization in the respiratory system of Drosophila. Developmental cell. 2008;15(4):617–26. doi: 10.1016/j.devcel.2008.08.017. PubMed PMID: 18854145; PubMed Central PMCID: PMCPMC2671232.

66. Fernandez-Capetillo O, Lee A, Nussenzweig M, Nussenzweig A. H2AX: the histone guardian of the genome. DNA repair. 2004;3(8-9):959–67. Epub 2004/07/29. doi: 10.1016/j.dnarep.2004.03.024. PubMed PMID: 15279782.

67. Le Guen T, Ragu S, Guirouilh-Barbat J, Lopez BS. Role of the double-strand break repair pathway in the maintenance of genomic stability. Mol Cell Oncol. 2015;2(1):e968020. doi: 10.4161/23723548.2014.968020. PubMed PMID: 27308383; PubMed Central PMCID: PMCPMC4905226.

68. Murr R. Interplay between different epigenetic modifications and mechanisms. Adv Genet. 2010;70:101–41. doi: 10.1016/B978-0-12-380866-0.60005-8. PubMed PMID: 20920747.

69. Madigan JP, Chotkowski HL, Glaser RL. DNA double-strand break-induced phosphorylation of Drosophila histone variant H2Av helps prevent radiation-induced apoptosis. Nucleic Acids Res. 2002;30(17):3698–705. PubMed PMID: 12202754; PubMed Central PMCID: PMCPMC137418.

70. Lake CM, Holsclaw JK, Bellendir SP, Sekelsky J, Hawley RS. The development of a monoclonal antibody recognizing the Drosophila melanogaster phosphorylated histone H2A variant (gamma-H2AV). G3. 2013;3(9):1539–43. doi: 10.1534/g3.113.006833. PubMed PMID: 23833215; PubMed Central PMCID: PMCPMC3755914.

71. Zhou F, Rasmussen A, Lee S, Agaisse H. The UPD3 cytokine couples environmental challenge and intestinal stem cell division through modulation of JAK/STAT signaling in the stem cell microenvironment. Developmental biology. 2013;373(2):383–93. Epub 2012/11/01. doi: 10.1016/j.ydbio.2012.10.023. PubMed PMID: 23110761; PubMed Central PMCID: PMC3534909.

72. Kasibhatla S, Amarante-Mendes GP, Finucane D, Brunner T, Bossy-Wetzel E, Green DR. Acridine Orange/Ethidium Bromide (AO/EB) Staining to Detect Apoptosis. CSH protocols. 2006;2006(3). Epub 2006/01/01. doi: 10.1101/pdb.prot4493. PubMed PMID: 22485874.

73. Goyal L, McCall K, Agapite J, Hartwieg E, Steller H. Induction of apoptosis by Drosophila reaper, hid and grim through inhibition of IAP function. The EMBO journal. 2000;19(4):589–97. doi: 10.1093/emboj/19.4.589. PubMed PMID: 10675328; PubMed Central PMCID: PMCPMC305597.

74. McEwen DG, Peifer M. Puckered, a Drosophila MAPK phosphatase, ensures cell viability by antagonizing JNK-induced apoptosis. Development. 2005;132(17):3935–46. doi: 10.1242/dev.01949. PubMed PMID: 16079158.

75. Ren F, Shi Q, Chen Y, Jiang A, Ip YT, Jiang H, et al. Drosophila Myc integrates multiple signaling pathways to regulate intestinal stem cell proliferation during midgut regeneration. Cell Res. 2013;23(9):1133–46. doi: 10.1038/cr.2013.101. PubMed PMID: 23896988; PubMed Central PMCID: PMCPMC3760623.

76. Shaw RL, Kohlmaier A, Polesello C, Veelken C, Edgar BA, Tapon N. The Hippo pathway regulates intestinal stem cell proliferation during Drosophila adult midgut regeneration. Development. 2010;137(24):4147–58. doi: 10.1242/dev.052506. PubMed PMID: 21068063; PubMed Central PMCID: PMCPMC2990206.

77. Amcheslavsky A, Jiang J, Ip YT. Tissue damage-induced intestinal stem cell division in Drosophila. Cell stem cell. 2009;4(1):49–61. Epub 2009/01/09. doi: 10.1016/j.stem.2008.10.016. PubMed PMID: 19128792; PubMed Central PMCID: PMC2659574.

78. Baker NE, Yu SY. The R8-photoreceptor equivalence group in Drosophila: fate choice precedes regulated Delta transcription and is independent of Notch gene dose. Mech Dev. 1998;74(1-2):3–14. PubMed PMID: 9651468.

79. Beebe K, Lee WC, Micchelli CA. JAK/STAT signaling coordinates stem cell proliferation and multilineage differentiation in the Drosophila intestinal stem cell lineage. Developmental biology. 2010;338(1):28–37. doi: 10.1016/j.ydbio.2009.10.045. PubMed PMID: 19896937.

80. Malumbres M, Barbacid M. Cell cycle, CDKs and cancer: a changing paradigm. Nature reviews Cancer. 2009;9(3):153–66. Epub 2009/02/25. doi: 10.1038/nrc2602. PubMed PMID: 19238148.

81. Nakamura M, Ohsawa S, Igaki T. Mitochondrial defects trigger proliferation of neighbouring cells via a senescence-associated secretory phenotype in Drosophila. Nature communications. 2014;5:5264. Epub 2014/10/28. doi: 10.1038/ncomms6264. PubMed PMID: 25345385.

82. Geisen C, Moroy T. The oncogenic activity of cyclin E is not confined to Cdk2 activation alone but relies on several other, distinct functions of the protein. The Journal of biological chemistry. 2002;277(42):39909–18. doi: 10.1074/jbc.M205919200. PubMed PMID: 12149264.

83. West CM, Dunning AM, Rosenstein BS. Genome-wide association studies and prediction of normal tissue toxicity. Semin Radiat Oncol. 2012;22(2):91–9. doi: 10.1016/j.semradonc.2011.12.007. PubMed PMID: 22385916.

84. Herskind C, Talbot CJ, Kerns SL, Veldwijk MR, Rosenstein BS, West CM. Radiogenomics: A systems biology approach to understanding genetic risk factors for radiotherapy toxicity? Cancer Lett. 2016;382(1):95– 109. doi: 10.1016/j.canlet.2016.02.035. PubMed PMID: 26944314; PubMed Central PMCID: PMCPMC5016239.

85. Borghini A, Vecoli C, Mercuri A, Petruzzelli MF, D’Errico MP, Portaluri M, et al. Genetic risk score and acute skin toxicity after breast radiation therapy. Cancer biotherapy & radiopharmaceuticals. 2014;29(7):267– 72. doi: 10.1089/cbr.2014.1620. PubMed PMID: 25099761.

86. Carulli AJ, Keeley TM, Demitrack ES, Chung J, Maillard I, Samuelson LC. Notch receptor regulation of intestinal stem cell homeostasis and crypt regeneration. Developmental biology. 2015;402(1):98–108. doi: 10.1016/j.ydbio.2015.03.012. PubMed PMID: 25835502; PubMed Central PMCID: PMC4433599.

87. Apidianakis Y, Rahme LG. Drosophila melanogaster as a model for human intestinal infection and pathology. Disease models & mechanisms. 2011;4(1):21–30. doi: 10.1242/dmm.003970. PubMed PMID: 21183483; PubMed Central PMCID: PMC3014343.

88. Mak KS, Gainor JF, Niemierko A, Oh KS, Willers H, Choi NC, et al. Significance of targeted therapy and genetic alterations in EGFR, ALK, or KRAS on survival in patients with non-small cell lung cancer treated with radiotherapy for brain metastases. Neurooncology. 2015;17(2):296–302. doi: 10.1093/neuonc/nou146. PubMed PMID: 25053852; PubMed Central PMCID: PMC4288518.

89. Kap EJ, Richter S, Rudolph A, Jansen L, Ulrich A, Hoffmeister M, et al. Genetic variants in the glutathione S-transferase genes and survival in colorectal cancer patients after chemotherapy and differences according to treatment with oxaliplatin. Pharmacogenetics and genomics. 2014;24(7):340–7. doi: 10.1097/FPC.0000000000000059. PubMed PMID: 24842074.

90. Pacelli R, Conson M, Cella L, Liuzzi R, Troncone G, Iorio V, et al. Radiation therapy following surgery for localized breast cancer: outcome prediction by classical prognostic factors and approximated genetic subtypes. Journal of radiation research. 2013;54(2):292–8. doi: 10.1093/jrr/rrs087. PubMed PMID: 23019151; PubMed Central PMCID: PMC3589925.

91. Freytag SO, Stricker H, Lu M, Elshaikh M, Aref I, Pradhan D, et al. Prospective randomized phase 2 trial of intensity modulated radiation therapy with or without oncolytic adenovirus-mediated cytotoxic gene therapy in intermediate-risk prostate cancer. International journal of radiation oncology, biology, physics. 2014;89(2):268–76. doi: 10.1016/j.ijrobp.2014.02.034. PubMed PMID: 24837889; PubMed Central PMCID: PMC4026796.

92. Bhatia S. Genetic variation as a modifier of association between therapeutic exposure and subsequent malignant neoplasms in cancer survivors. Cancer. 2015;121(5):648–63. doi: 10.1002/cncr.29096. PubMed PMID: 25355167; PubMed Central PMCID: PMC4339370.

93. Akasaka Y, Saikawa Y, Fujita K, Kubota T, Ishikawa Y, Fujimoto A, et al. Expression of a candidate marker for progenitor cells, Musashi-1, in the proliferative regions of human antrum and its decreased expression in intestinal metaplasia. Histopathology. 2005;47(4):348–56. doi: 10.1111/j.1365-2559.2005.02223.x. PubMed PMID: 16178889.

94. Potten CS, Booth C, Tudor GL, Booth D, Brady G, Hurley P, et al. Identification of a putative intestinal stem cell and early lineage marker; musashi-1. Differentiation. 2003;71(1):28–41. PubMed PMID: 12558601.

95. Kaneko Y, Sakakibara S, Imai T, Suzuki A, Nakamura Y, Sawamoto K, et al. Musashi1: an evolutionally conserved marker for CNS progenitor cells including neural stem cells. Dev Neurosci. 2000;22(1-2):139–53. doi: 10.1159/000017435. PubMed PMID: 10657706.

96. Phelps CB, Brand AH. Ectopic gene expression in Drosophila using GAL4 system. Methods. 1998;14(4):367–79. doi: 10.1006/meth.1998.0592. PubMed PMID: 9608508.

97. Okano H, Imai T, Okabe M. Musashi: a translational regulator of cell fate. J Cell Sci. 2002;115(Pt 7):1355–9. PubMed PMID: 11896183.

98. King RC, Darrow JB, Kaye NW. Studies on Different Classes of Mutations Induced by Radiation of Drosophila Melanogaster Females. Genetics. 1956;41(6):890–900. PubMed PMID: 17247670; PubMed Central PMCID: PMCPMC1224370.

99. Koval TM, Myser WC, Hart RW, Hink WF. Comparison of survival and unscheduled DNA synthesis between an insect and a mammalian cell line following X-ray treatments. Mutation research. 1978;49(3):431–5. PubMed PMID: 634308.

100. Ducoff HS. Causes of death in irradiated adult insects. Biol Rev Camb Philos Soc. 1972;47(2):211–40. PubMed PMID: 4206386.

101. Moskalev A, Zhikrivetskaya S, Krasnov G, Shaposhnikov M, Proshkina E, Borisoglebsky D, et al. A comparison of the transcriptome of Drosophila melanogaster in response to entomopathogenic fungus, ionizing radiation, starvation and cold shock. BMC Genomics. 2015;16 Suppl 13:S8. doi: 10.1186/1471-2164-16-S13-S8. PubMed PMID: 26694630; PubMed Central PMCID: PMCPMC4686790.

102. Xing Y, Su TT, Ruohola-Baker H. Tie-mediated signal from apoptotic cells protects stem cells in Drosophila melanogaster. Nature communications. 2015;6:7058. doi: 10.1038/ncomms8058. PubMed PMID: 25959206; PubMed Central PMCID: PMCPMC4451836.

103. Tricoire H, Rera M. A New, Discontinuous 2 Phases of Aging Model: Lessons from Drosophila melanogaster. PloS one. 2015;10(11):e0141920. doi: 10.1371/journal.pone.0141920. PubMed PMID: 26528826; PubMed Central PMCID: PMCPMC4631373.

104. Jiang H, Patel PH, Kohlmaier A, Grenley MO, McEwen DG, Edgar BA. Cytokine/Jak/Stat signaling mediates regeneration and homeostasis in the Drosophila midgut. Cell. 2009;137(7):1343–55. doi: 10.1016/j.cell.2009.05.014. PubMed PMID: 19563763; PubMed Central PMCID: PMC2753793.

105. Shlevkov E, Morata G. A dp53/JNK-dependant feedback amplification loop is essential for the apoptotic response to stress in Drosophila. Cell Death Differ. 2012;19(3):451–60. doi: 10.1038/cdd.2011.113. PubMed PMID: 21886179; PubMed Central PMCID: PMCPMC3278728.

106. Knoblich JA, Sauer K, Jones L, Richardson H, Saint R, Lehner CF. Cyclin E controls S phase progression and its down-regulation during Drosophila embryogenesis is required for the arrest of cell proliferation. Cell. 1994;77(1):107–20. PubMed PMID: 8156587.

107. Nejdfors P, Ekelund M, Westrom BR, Willen R, Jeppsson B. Intestinal permeability in humans is increased after radiation therapy. Diseases of the colon and rectum. 2000;43(11):1582–7; discussion 7-8. Epub 2000/11/23. PubMed PMID: 11089597.

108. Melichar B, Hyspler R, Kalabova H, Dvorak J, Ticha A, Zadak Z. Gastroduodenal, intestinal and colonic permeability during anticancer therapy. Hepato-gastroenterology. 2011;58(109):1193–9. Epub 2011/09/23. doi: 10.5754/hge08101. PubMed PMID: 21937377.

109. Vaisnav M, Xing C, Ku HC, Hwang D, Stojadinovic S, Pertsemlidis A, et al. Genome-wide association analysis of radiation resistance in Drosophila melanogaster. PloS one. 2014;9(8):e104858. doi: 10.1371/journal.pone.0104858. PubMed PMID: 25121966; PubMed Central PMCID: PMCPMC4133248.

110. Lanza A, Ravaud P, Riveros C, Dechartres A. Comparison of Estimates between Cohort and Case-Control Studies in Meta-Analyses of Therapeutic Interventions: A Meta-Epidemiological Study. PloS one. 2016;11(5):e0154877. doi: 10.1371/journal.pone.0154877. PubMed PMID: 27159025; PubMed Central PMCID: PMCPMC4861326.

111. de Araujo PR, Gorthi A, da Silva AE, Tonapi SS, Vo DT, Burns SC, et al. Musashi1 Impacts Radio-Resistance in Glioblastoma by Controlling DNA-Protein Kinase Catalytic Subunit. Am J Pathol. 2016;186(9):2271–8. doi: 10.1016/j.ajpath.2016.05.020. PubMed PMID: 27470713; PubMed Central PMCID: PMCPMC5012509.

112. Yousefi M, Li N, Nakauka-Ddamba A, Wang S, Davidow K, Schoenberger J, et al. Msi RNA-binding proteins control reserve intestinal stem cell quiescence. The Journal of cell biology. 2016;215(3):401–13. doi: 10.1083/jcb.201604119. PubMed PMID: 27799368; PubMed Central PMCID: PMCPMC5100293.

