## Supplementary figures and images for "*Musashi* expression in intestinal stem cells attenuates radiation-induced decline in intestinal homeostasis and survival in *Drosophila*"

### Supplemental Figure 1

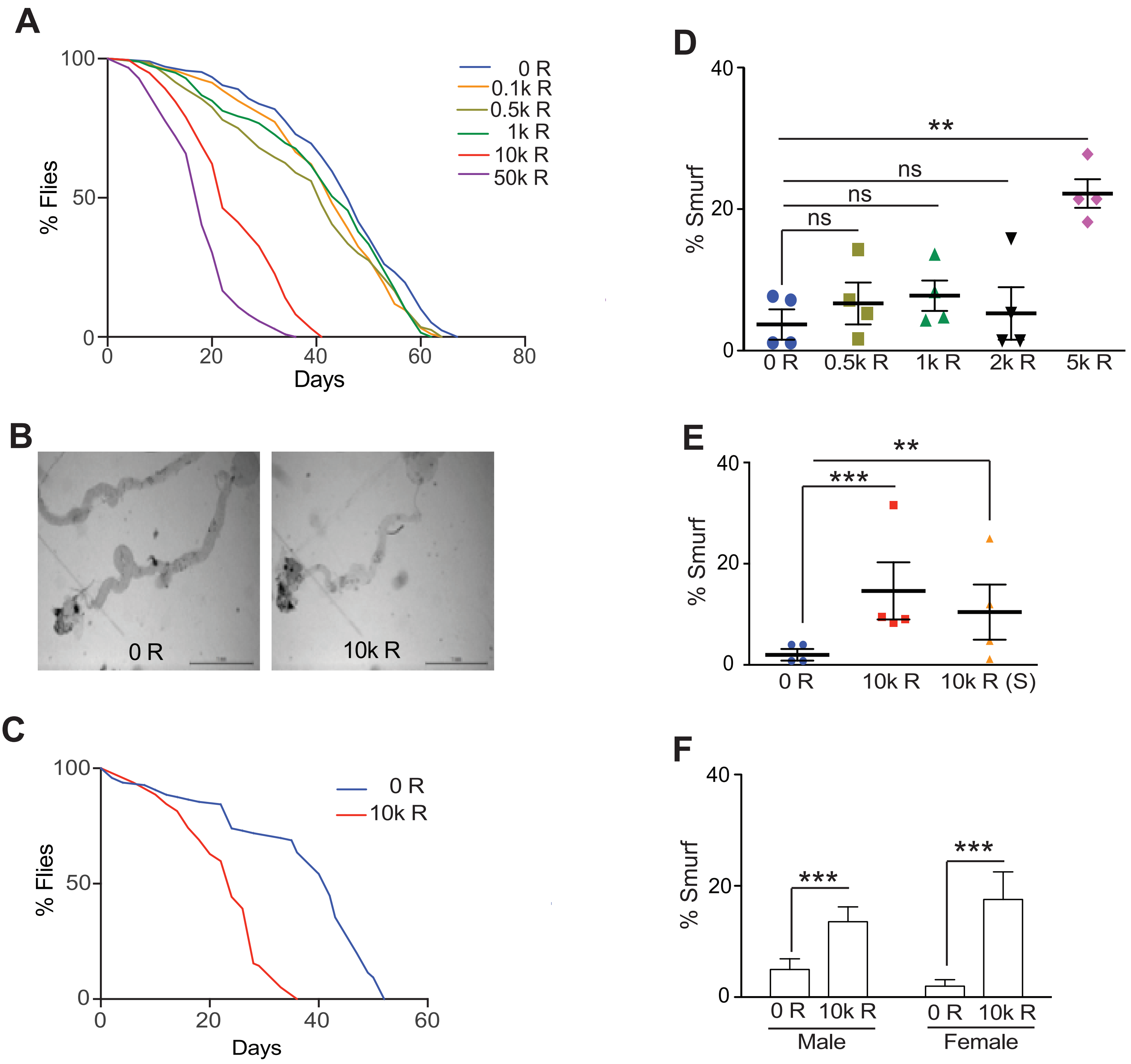

### Supplemental Figure 2

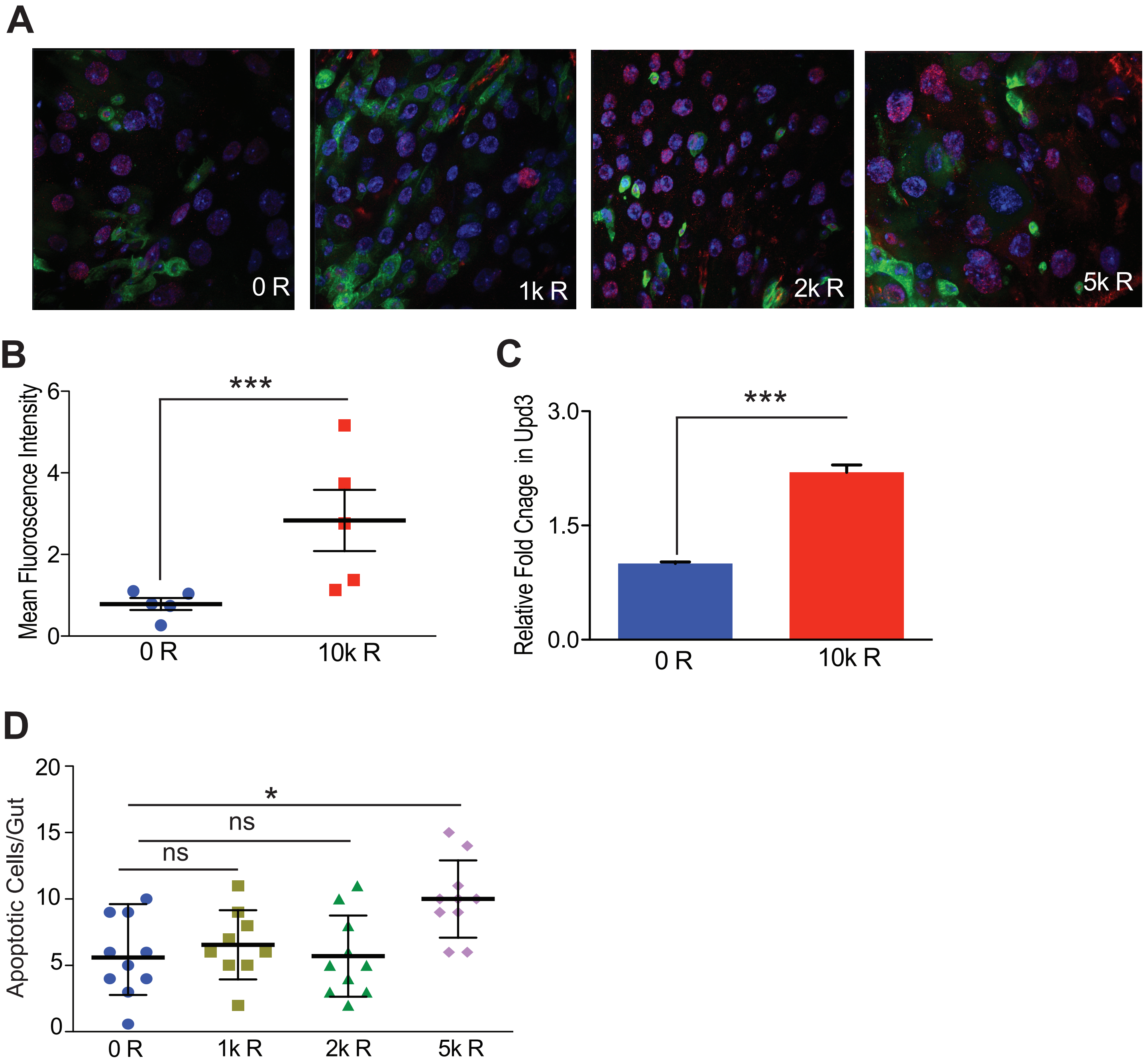

### Supplemental Figure 3

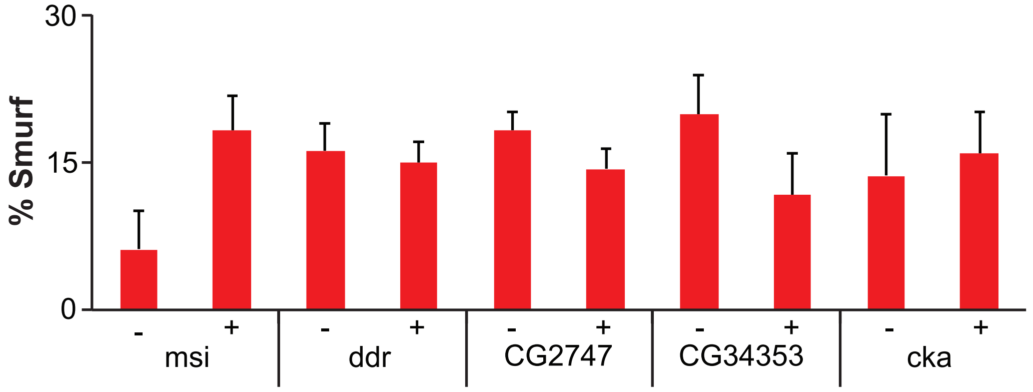
